# HiCanu: accurate assembly of segmental duplications, satellites, and allelic variants from high-fidelity long reads

**DOI:** 10.1101/2020.03.14.992248

**Authors:** Sergey Nurk, Brian P. Walenz, Arang Rhie, Mitchell R. Vollger, Glennis A. Logsdon, Robert Grothe, Karen H. Miga, Evan E. Eichler, Adam M. Phillippy, Sergey Koren

## Abstract

Complete and accurate genome assemblies form the basis of most downstream genomic analyses and are of critical importance. Recent genome assembly projects have relied on a combination of noisy long-read sequencing and accurate short-read sequencing, with the former offering greater assembly continuity and the latter providing higher consensus accuracy. The recently introduced PacBio HiFi sequencing technology bridges this divide by delivering long reads (>10 kbp) with high per-base accuracy (>99.9%). Here we present HiCanu, a significant modification of the Canu assembler designed to leverage the full potential of HiFi reads via homopolymer compression, overlap-based error correction, and aggressive false overlap filtering. We benchmark HiCanu with a focus on the recovery of haplotype diversity, major histocompatibility complex (MHC) variants, satellite DNAs, and segmental duplications. For diploid human genomes sequenced to 30× HiFi coverage, HiCanu achieved superior accuracy and allele recovery compared to the current state of the art. On the effectively haploid CHM13 human cell line, HiCanu achieved an NG50 contig size of 77 Mbp with a per-base consensus accuracy of 99.999% (QV50), surpassing recent assemblies of high-coverage, ultra-long Oxford Nanopore reads in terms of both accuracy and continuity. This HiCanu assembly correctly resolves 337 out of 341 validation BACs sampled from known segmental duplications and provides the first preliminary assemblies of 9 complete human centromeric regions. Although gaps and errors still remain within the most challenging regions of the genome, these results represent a significant advance towards the complete assembly of human genomes.

**Availability:** HiCanu is implemented within the Canu assembly framework and is available from https://github.com/marbl/canu.

## Introduction

Genome assembly is the process of reconstructing continuous genomic regions from shorter overlapping fragments, called reads (Nagarajan and Pop 2010; Miller et al. 2010). Recently, long-read sequencing technologies have significantly simplified assembly by generating multikilobase reads, which span most common genomic repeats (Koren et al. 2013; Chin et al. 2013; Koren and Phillippy 2014; Gordon et al. 2016; Bickhart et al. 2017; Kronenberg et al. 2018). Despite the per-base error rate of the input reads exceeding 10%, state of the art assembly methods are able to resolve instances of longer repeats with sequence divergence as low as 2% (Koren et al. 2017; Kolmogorov et al. 2019). However, a significant fraction of the human genome is represented by long segmental duplications of higher sequence identity. According to the current annotation of the human reference (Bailey et al. 2001, 2002), approximately 208 Mbp of sequence is contained within repeats longer than 20 kbp with sequence identity greater than 98%. Low accuracy of the long read technologies has also made continuous reconstruction of individual haplotypes very challenging since humans can average less than one heterozygous variant per 1 kbp. Typical assembly strategies collapse the genome first and phase afterwards by calling variants, partitioning the reads, and re-assembling (Chin et al. 2016; Seo et al. 2016). State-of-the-art methods integrate different sequencing technologies (Chaisson et al. 2019; Kronenberg et al. 2019) or parental information (Koren et al. 2018) to obtain chromosome-scale, haplotype-resolved assemblies. However, these approaches have the downside of collapsing multi-copy repeats in the assembly or not resolving alleles which differ at only a few positions.

Recently, Pacific Biosciences introduced a new data type, referred to as HiFi reads (Wenger et al. 2019). The process of generating HiFi reads involves DNA fragmentation; adapter ligation and fragment circularization; and multi-pass sequencing of the circularized fragments. The resulting signal is then computationally processed to obtain an accurate consensus sequence for each individual fragment. To ensure that each fragment undergoes sufficient sequencing passes to obtain a high consensus accuracy, HiFi sequencing libraries are size selected for a target fragment size (currently up to 25 kbp).

While the resulting read lengths are modest by the modern long read sequencing standards— PacBio CLR reads frequently exceed 50 kbp, and ultra-long Oxford Nanopore reads can exceed even 100 kbp (Jain et al. 2018b)—HiFi is a major leap forward in terms of long-read read accuracy. As the accuracy of other long-read technologies have not exceeded 95%, the median accuracy of current HiFi reads can exceed 99.9% (>Q30), making them a promising data type for separating highly similar repeat instances and alleles.

Early studies adopting HiFi sequencing demonstrated improved variant calling and repeat resolution (Wenger et al. 2019; Vollger et al. 2020). However, these early assemblies were limited to resolving repeats with greater than 1% sequence divergence, due to limitations of existing tools (Wenger et al. 2019). The recently developed Peregrine assembler (Chin and Khalak 2019) greatly reduced assembly runtime and improved consensus accuracy, removing the need for post-processing, but did not address the issue of suboptimal repeat resolution or allele separation. Other recent work combined HiFi sequencing with complementary data types, such as parental information (Wenger et al. 2019), Hi-C (Garg et al. 2019), and Strand-seq (Porubsky et al. 2019) to obtain chromosome-scale, haplotype-resolved assemblies.

In the following sections we present HiCanu, a modification of the Canu assembler (Koren et al. 2017) designed to take full advantage of the high accuracy of HiFi reads. By mitigating the remaining sequencing errors in the input data (including previously unreported recurrent sequence-specific errors), HiCanu is able to achieve the resolution of up to 99.99% identical genomic repeats. As a result, HiCanu surpasses both other HiFi and recent ultra-long Oxford Nanopore-based human assemblies in terms of both repeat resolution and per-base consensus accuracy. These prior assemblies have also required a final “polishing” step to improve consensus accuracy, which can introduce errors in repeat instances due to ambiguous read mappings (Miga et al. 2019). Due to the initial accuracy of HiFi reads, and the precise resolution of allelic variants and repeats, HiCanu assemblies do not require polishing. Furthermore, in contrast with most previous assemblers that implicitly ignore small heterozygous variants in a diploid genome, HiCanu accurately captures both alleles in large phase blocks. In particular, we demonstrate that HiCanu consistently recovers both haplotypes for the six canonical MHC typing genes in the human genome, improving upon recently developed HiFi-based methods for haplotype-resolved assembly (Garg et al. 2019; Porubsky et al. 2019).

## Results

### HiCanu overview

HiCanu builds on the original Canu pipeline, replacing or significantly modifying its core methods. Here we provide an overview of the new pipeline, highlighting the introduced changes, while a more detailed description of individual steps can be found in the Methods section. Whereas the original Canu pipeline starts with read self-correction, which can homogenize reads from different alleles or near-identical repeat instances, HiCanu begins by compressing all consecutive copies of the same nucleotide to a single base (e.g. “AA…” becomes “A”). In accordance with the earlier observation that misestimation of homopolymer length is the primary error mode of HiFi technology (Wenger et al. 2019), the resulting homopolymer-compressed reads (or “compressed reads*”* for short) accurately encode the transitions between different bases of the underlying genomic regions. The compressed reads are then trimmed based on their overlaps to other reads to remove any chimeric sequences or sequencing adapters (see Overlap Based Trimming in (Koren et al. 2017)), and the overlaps are recomputed on the trimmed reads. The Overlap Error Adjustment (OEA) module (Holt et al. 2002; Koren et al. 2017) examines read overlap pileups to identify remaining sequencing errors in the individual reads and recomputes overlap alignment identities. Following our observation that microsatellite repeat arrays are also prone to HiFi read errors, the OEA procedure was modified to ignore any differences within these regions when computing the final alignment identity of two overlapping reads. Together, homopolymer compression, pileup-based read correction, and ignoring differences in microsatellite repeats result in a drastic reduction of observed sequencing errors (Figure 1). Draft contigs are then formed from the adjusted overlaps using Canu’s Bogart module (Koren et al. 2017), modified to better handle heterozygous variants and consider only high-identity read overlaps. The final contig sequences are obtained by computing a consensus across the original, uncompressed reads, arranged according to the layout of their compressed versions. Similar to many modern assemblers, when faced with a diploid genome, HiCanu outputs contigs as “pseudo-haplotypes” that preserve local allelic phasing but may switch between haplotypes across longer distances. A single set of contigs representing all resolved alleles is output regardless of ploidy, and additional processing with a tool such as Purge_dups (Guan et al. 2020) is required to partition the contigs into primary and alternate allele sets.

**Figure 1.**
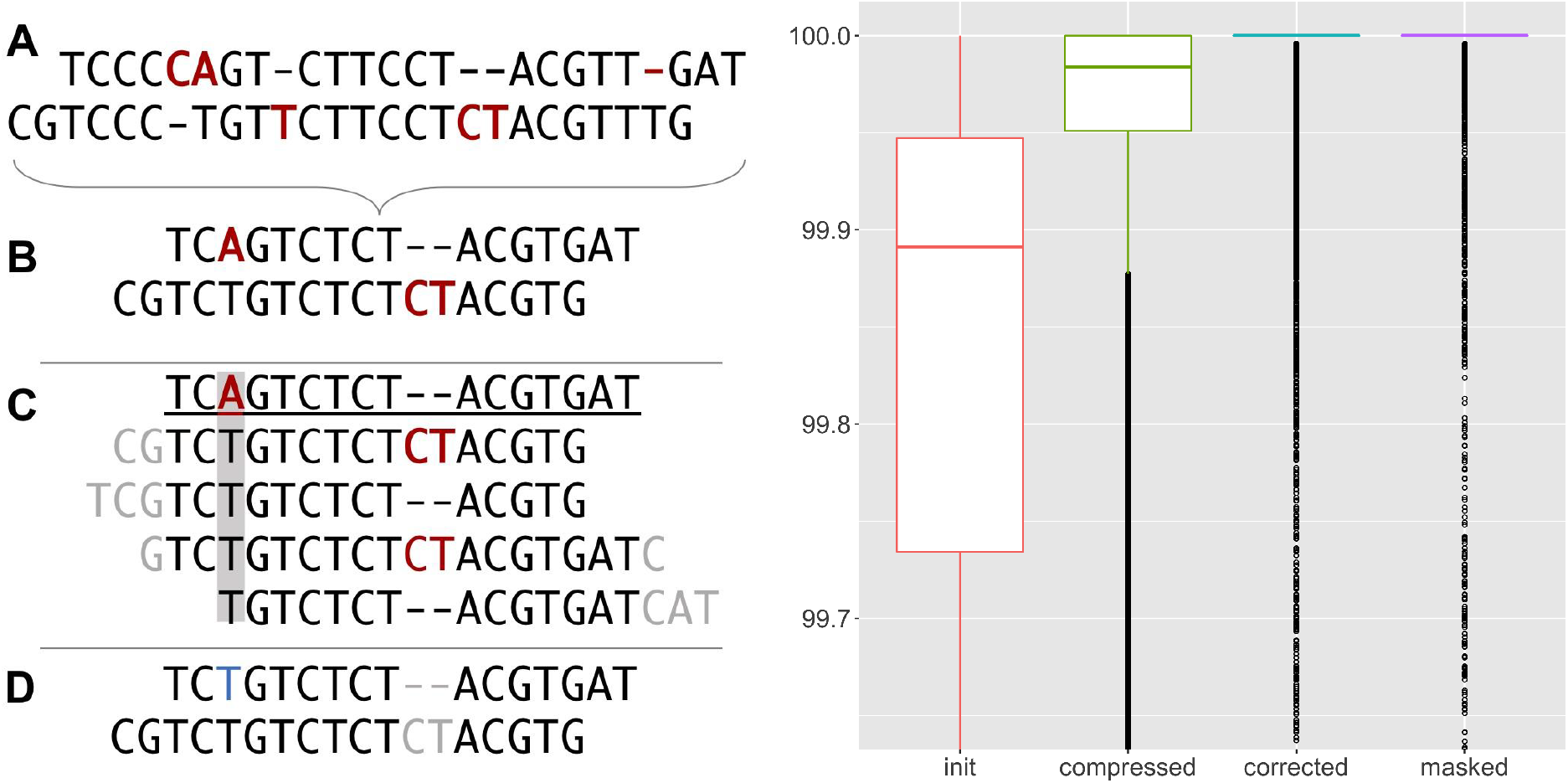
Impact of HiCanu processing on observed read quality. Left: A) Two hypothetical reads are shown with sequencing errors highlighted in red. B) The first step of HiCanu is to compress homopolymers, which obscures homopolymer length errors but retains enough information to accurately distinguish reads from different genomic loci. C) Overlaps are then computed for the compressed reads, and remaining errors are identified by examining the alignment pileups (gray rectangle). D) Finally, after correcting the identified errors (blue) and ignoring indels in regions of known systematic error (gray), the resulting overlap is 100% identical. Right: Sequence identity of reads from a 20 kbp HiFi library measured against the CHM13 chromosome X reference sequence v0.7 (Miga et al. 2019) after each step of HiCanu processing (Supplementary Note 1). Separate boxplots are shown for raw HiFi reads (init), homopolymer-compressed reads (compressed), OEA-corrected reads (corrected), and corrected reads after ignoring differences in microsatellite repeats (masked). The median read identity, indicated by solid segments, increases from less than 99.9% to 100% (note that the plots show an y-range of 99.65–100%). Supplementary Table 1 also shows how HiCanu processing increases the percentage of perfectly-aligned (100% identity) HiFi reads from less than 1% to over 97%.

### *Drosophila* genome assembly

We first evaluated HiCanu on a 24 kbp HiFi library from a *Drosophila melanogaster* F1 hybrid (ISO1×A4) (Data Availability). To match typical coverage, the HiFi dataset was downsampled to 40× and assembled with the HiFi-specific tools, HiCanu and Peregrine (Chin and Khalak 2019), as well as the conventional long-read assembler, Canu. Canu was chosen as it was previously shown to achieve the highest assembly continuity and superior repeat resolution among other popular long-read assemblers on HiFi data (Wenger et al. 2019). For comparison, we also include a Canu assembly of 200× PacBio single-pass reads (CLR) for the same organism. Contigs less than 50 kbp were filtered from the assemblies in order to exclude low-quality sequences consisting of only a few reads.

Total assembly size varied between HiCanu (301 Mbp), Canu (293 Mbp), Peregrine (162 Mbp), and CLR (294 Mbp). Besides Peregrine, the assembly sizes were more than twice that of the 144 Mbp *D. melanogaster* haploid reference genome (Hoskins et al. 2015), suggesting that both haplotypes of the highly heterozygous F1 were successfully assembled (heterozygosity estimated at 0.7% by Genomescope (Vurture et al. 2017), Supplementary Figure 1). The large fraction of duplicated BUSCO (Waterhouse et al. 2018) genes also supported the hypothesis that the assemblies captured alleles from both haplotypes (Supplementary Table 2). To facilitate like-for-like comparison of all assemblies, we ran Purge_dups (Guan et al. 2020) to identify alleles and divide the assemblies into primary and alternate contig sets (Methods). Assembly statistics were then computed for both contig sets and the results summarized in Table 1. Perbase consensus accuracy was estimated using Merqury (Rhie et al. 2020) with Illumina sequencing data from the *D. melanogaster* F1 parental strains (Supplementary Note 2).

**Table 1.**
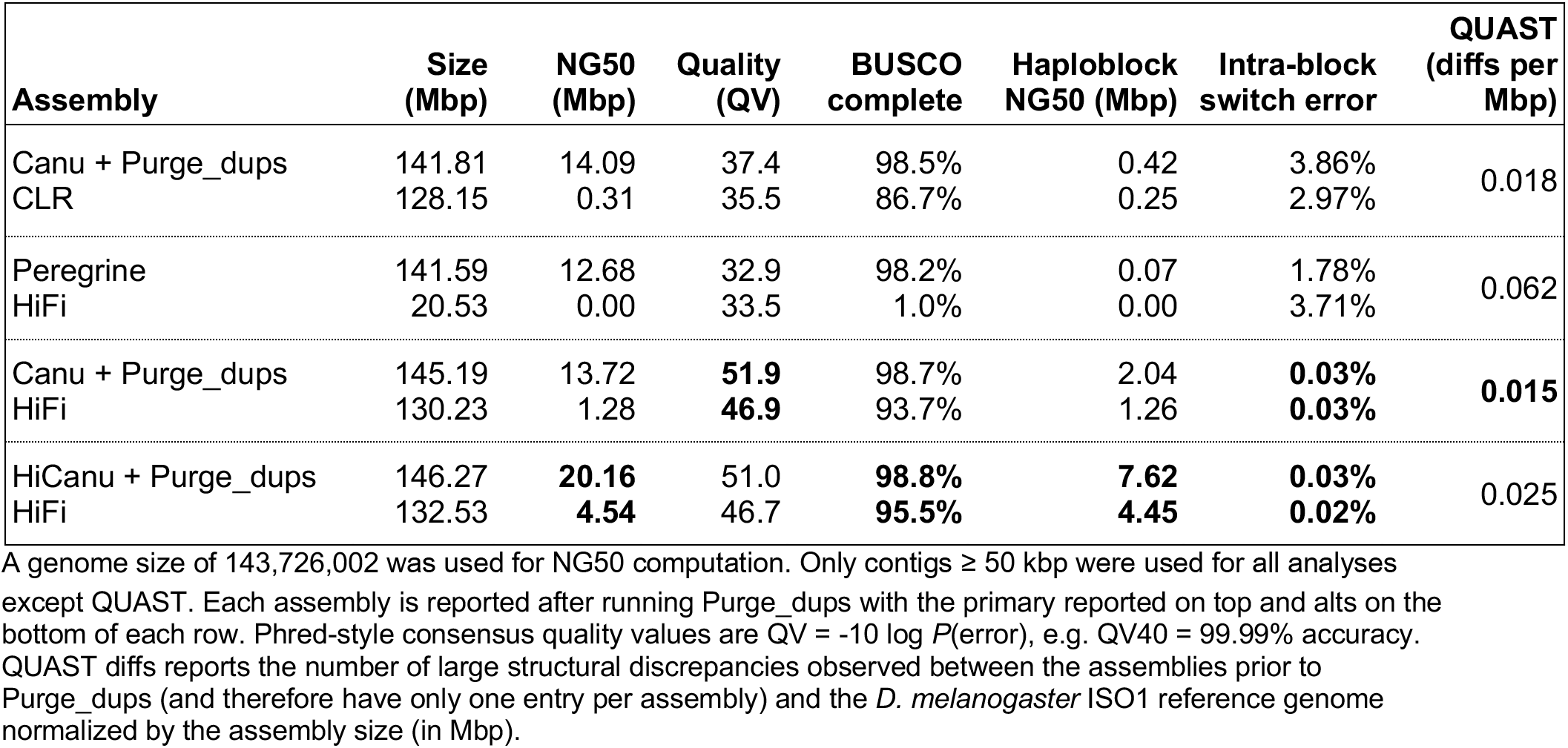
*D. melanogaster* ISO1×A4 assembly benchmarking results for PacBio CLR and HiFi.

The primary contig sets across all assemblies reported high BUSCO completeness (>98%). BUSCO duplication values were <2% across all contig sets. The HiCanu primary contig set was noticeably more continuous than any other assembly as measured by NG50 (N such that half the haploid genome size is represented by contigs of this size or greater). Canu and HiCanu showed very similar per-base consensus accuracy, radically improving on both Peregrine and CLR assemblies. The Peregrine assembly collapsed both haplotypes together and output few alternate contigs (total length <21 Mbp). HiCanu improved over all other assemblies with respect to the total size, BUSCO completeness, and continuity of the alternate set (including a 3-fold improvement in NG50 over Canu).

To assess the integrity of the assemblies we used QUAST v5.02 (Gurevich et al. 2013) to compare the assemblies against the chromosome arms of the *D. melanogaster* ISO1 reference. Since Purge_dups can split and/or trim the initial contigs, but has a negligible effect on continuity, we report structural correctness of the original assemblies. Considering that one of the haplotypes (derived from the A4 parental strain) is expected to differ significantly from the reference, we adjusted QUAST’s parameters to detect only large scale genomic differences (Methods). While the HiCanu assembly reported three more structural discrepancies than Canu (7 vs 4), it maintained the highest NG50 and alternate contig BUSCO completeness.

In general, HiFi reads alone cannot be used to infer phasing across homozygous regions longer than the read length, so the contigs produced by HiCanu (and Canu) represent “pseudohaplotypes”, which may switch between haplotypes. However, for highly heterozygous genomes with short regions of homozygosity, HiCanu is expected to produce a low number of haplotype switches and mostly preserve long-range phasing. We used Merqury (Rhie et al. 2020) to split the initial contigs into continuous phase blocks, based on haplotype-specific k-mer markers inferred from parental Illumina reads (Supplementary Note 2). As a baseline, we considered a haplotype-resolved assembly produced by TrioCanu (Koren et al. 2018) generated using a combination of CLR reads and parental Illumina data. The HiCanu primary (alternate) contig set has an estimated phase block NG50 of 7.62 Mbp (4.45 Mbp), a maximal block length of 25.4 Mbp (10.1 Mbp), and a low percentage of discordant markers within predicted haplotype blocks (switch rate) of 0.03% (0.02%). For comparison, the TrioCanu assembly, has a paternal-ISO (maternal-A4) phase block NG50 of 13.9 Mbp (21.39 Mbp), a max block size of 24.7 Mbp (27.7 Mbp), and an intra-block switch rate of 0.1% (0.04%). In contrast, the phase blocks of all other considered assemblies are much less continuous (at least a 3.5-fold drop in phase block NG50 compared to HiCanu) and, in the case of Peregrine and CLR assemblies, a much higher switch error (Table 1).

### Human genome assemblies

We next ran HiCanu, Canu, and Peregrine on three different human datasets (Data Availability): a 20 kbp library of the completely homozygous cell line CHM13 (Kronenberg et al. 2018; Miga et al. 2019; Vollger et al. 2020), a 15 kbp library of the Ashkenazic cell line HG002 from the Personal Genome Project (Church 2005; Wenger et al. 2019), and a combined library (12% 10 kbp, 62% 15 kbp, 26% 20 kbp) for the Puerto Rican cell line HG00733 from the 1000 Genomes Project (1000 Genomes Project Consortium et al. 2012; Porubsky et al. 2019). All datasets consist of approximately 30× HiFi sequencing coverage. For the HG002 dataset, we re-used the best assembly from a recent study (Wenger et al. 2019) as it reflects a Canu 1.7.1 assembly prior to HiCanu’s development and the associated improvements to Canu’s core modules. We additionally included the most continuous published (pseudo-haplotype) assemblies of the same genomes, which relied on ultra-long Oxford Nanopore (ONT) reads to achieve state-of-the-art repeat resolution (Shafin et al. 2019; Miga et al. 2019). As before, contigs less than 50 kbp were excluded from analysis. As the sizes of HiCanu assemblies for the diploid datasets HG002 and HG00733 were 5.30 Gbp and 5.46 Gbp, respectively, compared to a haploid genome size of 3.2 Gbp, we again post-processed all diploid assemblies with Purge_dups and computed statistics for both primary and alternate contig sets.

Per-base consensus quality was estimated by Merqury (Rhie et al. 2020) using Illumina data from the corresponding genome (Supplementary Note 2). To assess the structural correctness of the assemblies we followed the methodology of (Shafin et al. 2019). Namely, structural differences reported by QUAST v5.0.2 versus the human reference genome GRCh38 (Schneider et al. 2017) were post-processed to ignore breakpoints in centromeric regions and annotated segmental duplications, in order to reduce the number of false positives (Methods, Supplementary Table 3). As before, since Purge_dups may introduce or correct mis-assemblies as it modifies the contigs, the structural correctness assessment was performed on the original assemblies.

Primary contig summary statistics for the three human genomes are presented in Table 2. The continuity of HiCanu assemblies, as measured by NG50, exceeded that of all other HiFi-based assemblies and even rivaled the continuity of Nanopore ultra-long read assemblies. Reported rates of structural differences for HiCanu was on par with the other assemblies. For consensus accuracy, the HiCanu primary contig sets exceeded QV50 (99.999% accuracy) and alternate contigs sets exceeded QV40 (99.99% accuracy), while the unpolished Nanopore assemblies failed to exceed QV30 (99.9%). Although Nanopore assemblies currently require polishing with complementary technologies to maximize consensus accuracy, we discourage polishing HiCanu HiFi assemblies, because the available polishing pipelines may map reads back to the wrong repeat copies and actually introduce errors during polishing.

**Table 2.**
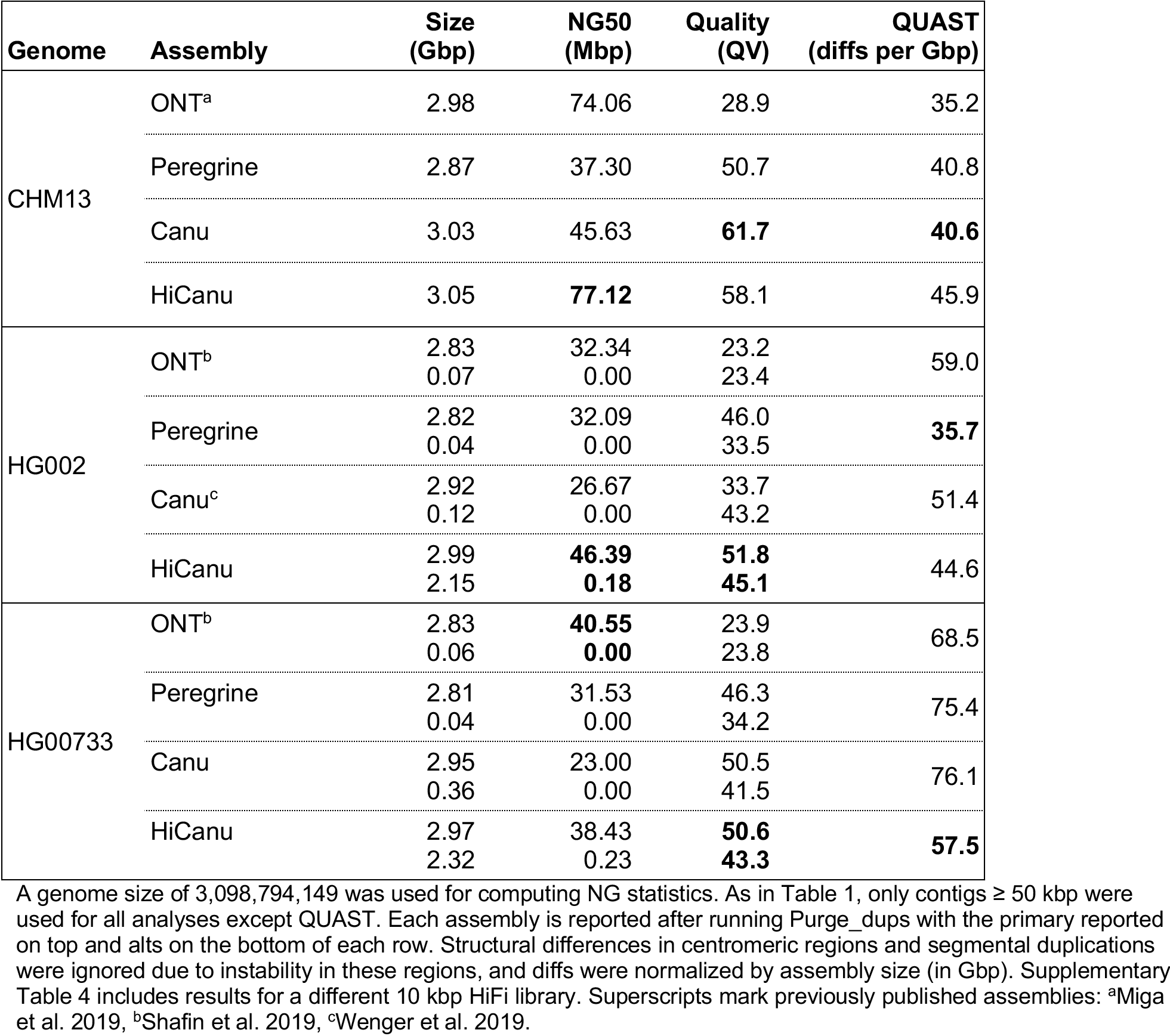
Human assembly benchmarking results for ultra-long Nanopore (ONT) and PacBio HiFi.

| Genome | Assembly | Size (Gbp) | NG50 (Mbp) | Quality (QV) | QUAST (diffs per Gbp) |
| --- | --- | --- | --- | --- | --- |
| CHM13 | ONT<sup>a</sup> | 2.98 | 74.06 | 28.9 | 35.2 |
|  | Peregrine | 2.87 | 37.30 | 50.7 | 40.8 |
|  | Canu | 3.03 | 45.63 | **61.7** | **40.6** |
|  | HiCanu | 3.05 | **77.12** | 58.1 | 45.9 |
| HG002 | ONT<sup>b</sup> | 2.83<br>0.07 | 32.34<br>0.00 | 23.2<br>23.4 | 59.0 |
|  | Peregrine | 2.82<br>0.04 | 32.09<br>0.00 | 46.0<br>33.5 | **35.7** |
|  | Canu<sup>c</sup> | 2.92<br>0.12 | 26.67<br>0.00 | 33.7<br>43.2 | 51.4 |
|  | HiCanu | 2.99<br>2.15 | **46.39**<br>**0.18** | **51.8**<br>**45.1** | 44.6 |
| HG00733 | ONT<sup>b</sup> | 2.83<br>0.06 | **40.55**<br>**0.00** | 23.9<br>23.8 | 68.5 |
|  | Peregrine | 2.81<br>0.04 | 31.53<br>0.00 | 46.3<br>34.2 | 75.4 |
|  | Canu | 2.95<br>0.36 | 23.00<br>0.00 | 50.5<br>41.5 | 76.1 |
|  | HiCanu | 2.97<br>2.32 | 38.43<br>0.23 | **50.6**<br>**43.3** | **57.5** |

Total length of the HiCanu alternate contig sets exceeded 2 Gbp, highlighting its ability to separate human alleles (corresponding values across other assemblies did not exceed 400 Mbp). The following section “Human haplotype phasing” further explores allele separation and phasing across these assemblies. The drastic improvements in consensus accuracy and allele separation for Canu versus HiCanu assemblies of HG002 is likely due to Canu improvements and bug fixes made during the HiCanu development process, whereas the CHM13 and HG00733 assemblies represent the latest Canu version and the differences are less pronounced.

For CHM13 and HG00733 genomes, we additionally validated the assemblies against long continuous fragments of the corresponding genome, earlier reconstructed via bacterial artificial chromosome (BAC) sequencing (Data Availability, no BACs were available for HG002). Many of these so-called “challenge” BACs were deliberately selected from genomic regions which pose significant assembly challenges (i.e. regions with segmental duplications), making them useful for assembly benchmarking (Chin and Khalak 2019; Shafin et al. 2019; Miga et al. 2019; Vollger et al. 2020). Table 3 summarizes how well the challenge BACs are captured by different assemblies. To recognize a BAC as “resolved” within the assembly, we required 99.5% of the BAC length to be aligned to a single contig by Minimap2 (Li 2018) (Methods). Note that HiCanu resolved the highest number of BACs across all considered assemblies and also achieved the highest BAC alignment quality (Table 3).

**Table 3.** “Challenge” BAC validation of human assemblies.

| Genome | Assembly | #BACs Resolved | Median QV |
| --- | --- | --- | --- |
| CHM13 | ONT<sup>a</sup> | 314/341 | 23.3 |
|  | Peregrine | 136/341 | 37.3 |
|  | Canu | 308/341 | 40.6 |
|  | HiCanu | **326/341** | **40.7** |
| HG00733 | ONT<sup>b</sup> | 124/179 | 18.9 |
|  | Peregrine | 74/179 | 27.0 |
|  | Canu | 122/179 | 28.2 |
|  | HiCanu | **164/179** | **33.9** |

The criteria for considering a BAC “resolved” is described in the main text and Methods. The alignment identity of each resolved BAC was computed individually and the median of these values reported as a Phred-style quality value. No validation BACs were available for HG002. Superscripts mark previously published assemblies: <sup>a</sup>Miga et al. 2019), <sup>b</sup>Shafin et al. 2019.

A deeper investigation of the unresolved CHM13 BAC sequences indicated that 11 BACs likely contain assembly errors or cloning artifacts themselves (Supplementary Note 3 and Supplementary Figures 2–5). Manual inspection of HiFi read alignments did not reveal any standard mis-assembly signatures in the corresponding regions of the HiCanu assembly, providing evidence that HiCanu was correct in these cases and able to resolve 337 out of 341 (99%) of the CHM13 challenge BACs (Supplementary Table 5).

While the challenge BACs are useful for validation, they do not represent the full landscape of human repeats. To further assess the ability of HiFi reads and different assemblers to resolve genomic repeats, we used the method of (Vollger et al. 2019) to identify collapsed repeat instances in the CHM13 assemblies. We identified approximately 21.7 Mbp of collapsed repeats corresponding to at least 56 Mbp of unresolved repetitive sequence. The HiCanu assembly had the lowest number of bases in regions identified as collapsed repeats, and the smallest amount of repetitive sequence predicted to be missing from the assembly (Supplementary Table 6, Supplementary Figure 6). A complementary mapping-based analysis confirmed the comparatively high completeness of the HiCanu assembly and classified the majority (80%) of missing sequence as satellite repeats, suggesting good recovery of all other human repeat classes (Supplementary Figure 6).

#### Human haplotype phasing

When assembling a diploid genome, an assembler must choose to either collapse alleles into a single sequence or preserve them as two separate sequences. Collapsing heterozygosity results in a mosaic consensus that may not faithfully represent any allele and can introduce frameshifting errors within coding sequence.

HiCanu assemblies of the diploid human genomes included more than 2 Gbp of alternate contigs, with high BUSCO completeness for both primary and alternate contig sets (>94% and >75%, respectively, Supplementary Table 7). We again used Merqury (Rhie et al. 2020) to analyze the phase blocks using parental Illumina data (Supplementary Note 2). The phase block NG50s of HiCanu primary (0.6 Mbp) and alternate (0.1 Mbp) contig sets were the highest across all considered assemblies (2.5-fold higher than next best, Supplementary Table 7). Note that the human phase block NG50s are significantly shorter than for the *D. melanogaster* F1 hybrid, but are longer than a typical human gene. For comparison, Supplementary Table 7 also includes statistics for the recently obtained haplotype-resolved assemblies of HG002 (Garg et al. 2019) and HG00733 (Porubsky et al. 2019). These recent studies have shown that multimegabase NG50 phase blocks can be obtained by integrating HiFi data with long-range linking information derived from Hi-C (Garg et al. 2019) or Strand-seq data (Porubsky et al. 2019).

To assess the ability of different tools to accurately recover complex, clinically relevant alleles, we compared assembly typing results for the six classical human leukocyte antigen (HLA) genes (Dilthey et al. 2019) to the known alleles for HG002 and HG00733, obtained by previous studies (Chin et al. 2019; Shafin et al. 2019) (Supplementary Table 8). Only HiCanu and TrioCanu were able to recover all alleles with 100% sequence identity (Supplementary Tables 8 and 9). The HiCanu contigs expectedly switch between the haplotypes, but there is only one switch in the MHC region. The Hi-C-phased HG002 assembly from (Garg et al. 2019) is phased across the length of the MHC region but contains consensus errors (e.g. both HLA-DRB1 alleles). The Strand-seq-phased HG00733 assembly from (Porubsky et al. 2019) is also phased across the length of the MHC region but incorrectly represents HLA-A and HLA-B as homozygous (with both alleles in the assembly matching one ground-truth allele). Both the Garg et al. and Porubsky et al. methods rely on initially collapsed assemblies that are then phased using the long-range data. These results suggest that separation of haplotypes early in the assembly process (rather than trying to recover them from collapsed assemblies) may improve the accurate recovery of heterozygous variation.

#### Complex regions of the CHM13 human genome

The CHM13 HiCanu assembly (Tables 2, 3, Figure 2) exceeded the predictions of our prior model of human assembly continuity (Supplementary Note 5, Supplementary Figure 7). To validate this result, we focused on the performance of HiCanu within some of the most difficult-to-assemble regions of the genome, namely centromeres and segmental duplications. Unlike past assemblies of the human genome, including clone-based assemblies, HiCanu generated several contigs spanning megabases of satellite DNA. The CHM13 HiCanu assembly contains 9 of 23 (39%) expected centromere regions; chromosomes 2, 3, 7, 8, 10, 12, 16, 19, and 20 (Supplementary Note 6, Supplementary Table 10). The structure of these regions was consistent with an expectation of one or more higher-order repeat (HOR) array(s) flanked by more divergent tracts of monomeric satellite DNA (Willard and Waye 1987; Schueler et al. 2001; She et al. 2004). Mapped read depth across these contigs shows relatively uniform sequence coverage that spans the α-satellite HOR array(s) into the unique sequences on the p- and q-arms (Figure 3, Supplementary Note 6, Supplementary Figure 8). The structure and length of the centromeric HOR array(s) in each contig is highly concordant with those reported in the literature (reviewed in McNulty and Sullivan 2018).

It is noteworthy that HiCanu generated a draft assembly of the CHM13 chromosome 19 centromere (Figure 3). Historically, this region has been considered to be one of the more challenging centromeres to reconstruct, since it carries multiple HOR tracts and shares α-satellite sequences with the centromere regions from chromosomes 1 and 5 (Hulsebos et al. 1988; Baldini et al. 1989; Pironon et al. 2010; Sullivan et al. 2017; McNulty and Sullivan 2018). HiCanu was not only able to assemble a contig that completely spans this centromere but also accurately distinguished three distinct HOR tracts (D19Z1, D19Z2, and D19Z3) (Supplementary Note 6, Supplementary Figure 9). This contig revealed a more complete representation of the HOR structure of the D19Z1 HOR unit (13-mer vs. 10-mer, Supplementary Figure 9a, Supplementary Figure 10) (Hulsebos et al. 1988; Puechberty et al. 1999), a chromosome 19-specific dimeric HOR (D19Z3, Supplementary Figure 9b, Supplementary Figure 10) (Baldini et al. 1989; Finelli et al. 1996), and two complex HORs (expected to represent D19Z2, Supplementary Note 6, Supplementary Figure 10). Alignment of HiFi sequence data to the corresponding HiCanu contig did not reveal any coverage anomalies (e.g. large dips or spikes) that could indicate the presence of structural errors. However, marker-assisted alignment of ultra-long Oxford Nanopore data (Miga et al. 2019), an orthogonal dataset, showed a coverage drop coinciding with the D19Z1 array. This may indicate a mis-assembly, mis-mapping of the noisy sequencing data, or biases in sequencing coverage. Due to the lack of a validated truth set in such regions, this will require extensive wet-lab validation and is left for future work.

**Figure 2.**
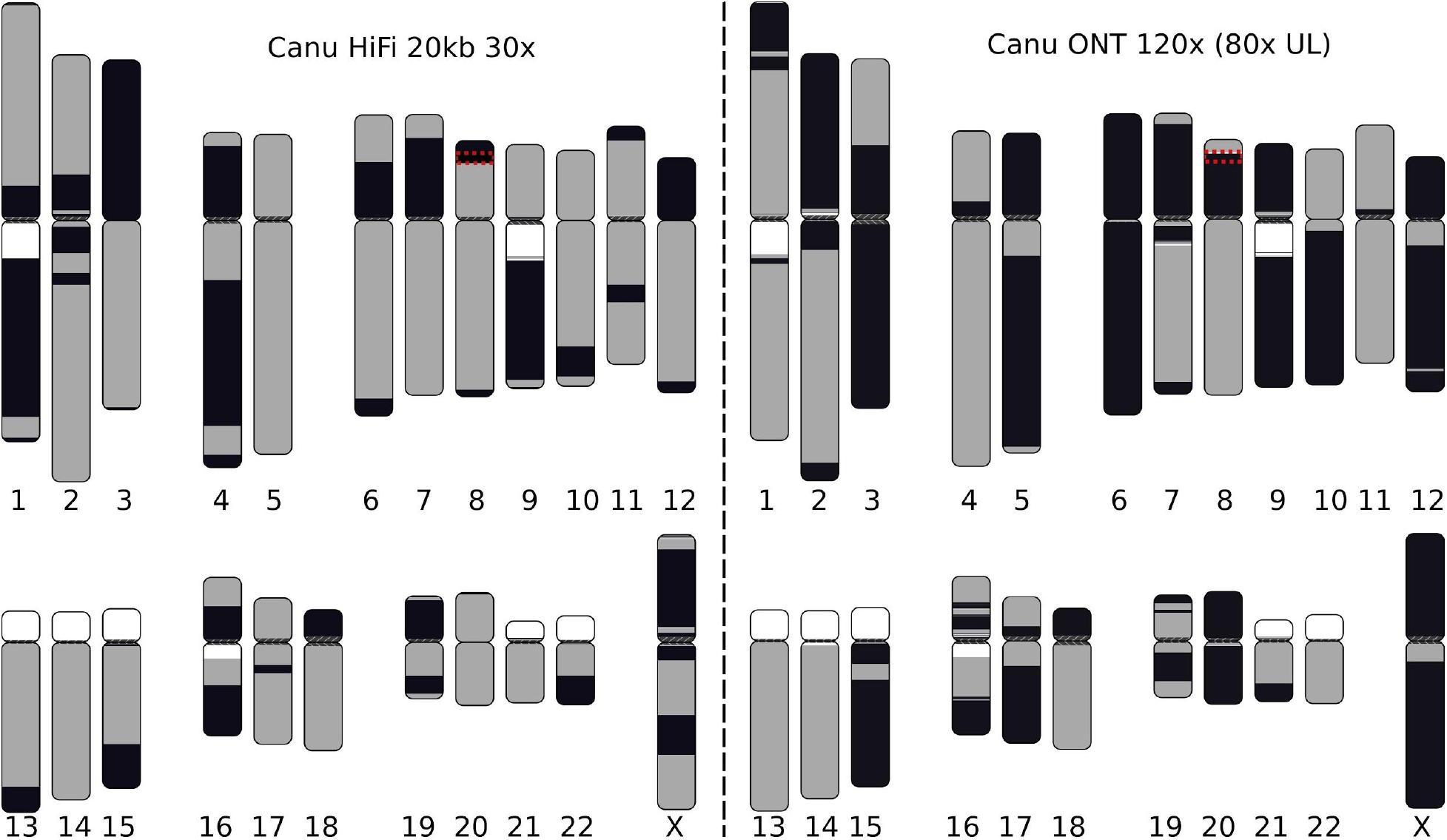
Visual representation of the most continuous HiFi-based and Nanopore-based assemblies of the CHM13 genome. HiCanu assembly of the 20 kbp HiFi dataset (left) and Canu assembly of an ultra-long Nanopore dataset (right). White regions indicate gaps in the current reference genome, while each gray and black block indicates a continuous contig alignment. Color switches from gray to black represent either the end of a contig or an alignment break. Assemblies were aligned to GRCh38 using MashMap (Jain et al. 2018a) and plots were generated using coloredChromosomes (Böhringer et al. 2002) as previously described (Berlin et al. 2015; Jain et al. 2018b). Note that some chromosomes (e.g. chrX) are better resolved by the Nanopore assembly due to the presence of near-perfect repeats. At the same time, chromosomes containing more diverged repeats (e.g. chr7 and chr16) are better resolved by the HiFi assembly. We note that some gaps in the HiFi assembly are caused by sequence-specific biases of current HiFi sequencing protocols (Supplementary Note 4). The red box highlights the β-defensin gene cluster on chromosome 8 which is split in both assemblies and detailed in Figure 4.

**Figure 3.**
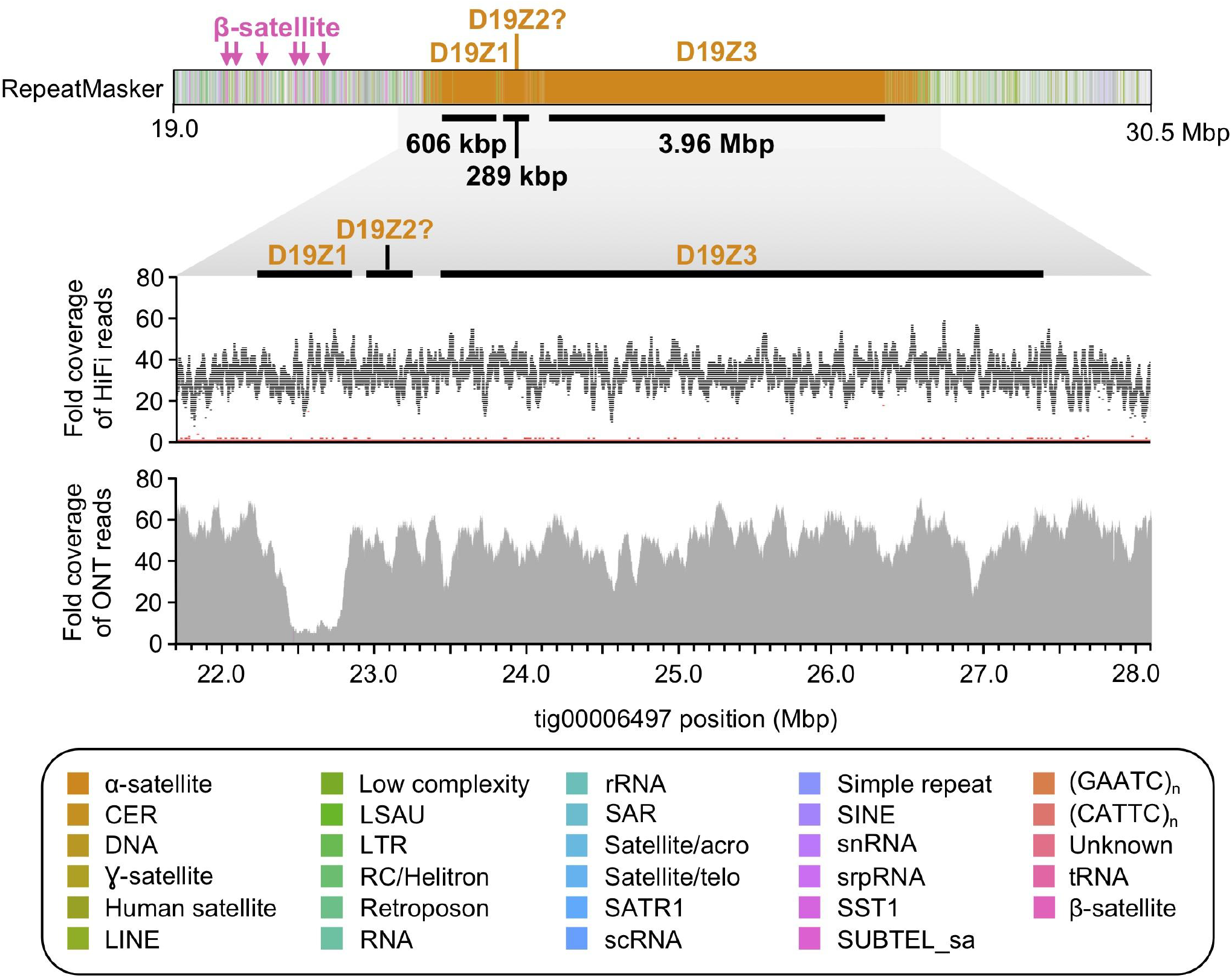
HiCanu assembly of the CHM13 chromosome 19 centromere. RepeatMasker (Smit et al. 2013) of tig00006497 reveals three α-satellite HOR arrays that reside within the chromosome 19 centromere (D19Z1, D19Z2?, and D19Z3; marked with black bars). These HOR arrays are 606 kbp, 289 kbp, and 3.96 Mbp in length, respectively, and are composed of a 13-mer, a complex higher-order HOR, and a dimeric HOR unit, respectively. The HOR repeat underlying D19Z2 shares limited sequence identity with the pG-A16 repeat previously described (Hulsebos et al. 1988; Choo et al. 1991; Finelli et al. 1996) and, therefore, is designated with a question mark. The α-satellite HOR arrays have relatively uniform coverage of HiFi and ultra-long Oxford Nanopore data, except for a drop in Oxford Nanopore sequencing coverage over the D19Z1 array, which may be due to a mis-assembly, read mis-mapping, or biases in sequencing. The HiFi coverage plot shows fold coverage of the most common base (black) and the second most common base (red).

Beyond the obvious challenge of centromere assembly, segmental duplications (SDs) represent another significant impediment and have been estimated to account for 68% of mis-assemblies and contig breaks in recent long-read genome assemblies, irrespective of the platform or assembly algorithm (Porubsky et al. 2019). To estimate the effect of SDs on the continuity of HiCanu assemblies, we aligned contigs from the CHM13 genome assemblies to the human reference genome (GRCh38) and tested if the ends of contigs mapped within SDs. Compared to Canu, Peregrine, or ONT assemblies, HiCanu had the fewest contig breaks within SDs (n=95) and the smallest overall fraction of contig ends mapping to SDs (49%) (Table 4). Of these 95 regions, 59 (62%) correspond to the longest (>10 kbp), most identical (> 98%), copy-number polymorphic duplicated regions of the human genome (Supplementary Figure 11). These results indicate that SDs are better resolved using HiCanu; however, SDs still contribute disproportionately to the overall number of assembly breaks.

**Table 4.** CHM13 contig ends found within segmental duplications.

| **Assembly** | **Total # of aligned contig ends** | **Total # of contig ends within SDs** | **Percent of contig ends within SDs** |
| --- | --- | --- | --- |
| ONT<sup>a</sup> | 202 | 137 | 68% |
| Canu | 322 | 170 | 53% |
| Peregrine | 468 | 398 | 85% |
| HiCanu | **192** | **95** | **49%** |

Previously published assemblies: <sup>a</sup>Miga et al. 2019.

The β-defensin gene cluster on chromosome 8 is a case in point. This approximately 6 Mbp region plays an important role in immune function and disease (Weinberg et al. 2012; Mohajeri et al. 2016) and is known to be highly repetitive and difficult to assemble (Bakar et al. 2009). Previous reconstructions have relied on a BAC-by-BAC assembly approach (Mohajeri et al. 2016), and the first continuous assembly of this region in CHM13 was obtained via manual assembly of both HiFi and ultra-long Nanopore data (Logsdon *et al*. in prep). Figure 4 illustrates self-alignment dot plots of the defensin region from the T2T chromosome 8 v3.0 assembly (Data Availability), as well as the *de novo* assembled contigs mapped against it. Both Canu and HiCanu assemblies of the HiFi data consist of four contigs without structural errors. In contrast, the complex inverted repeat structures resulted in mis-assembled and fragmented contigs in all other evaluated assemblies.

**Figure 4.**
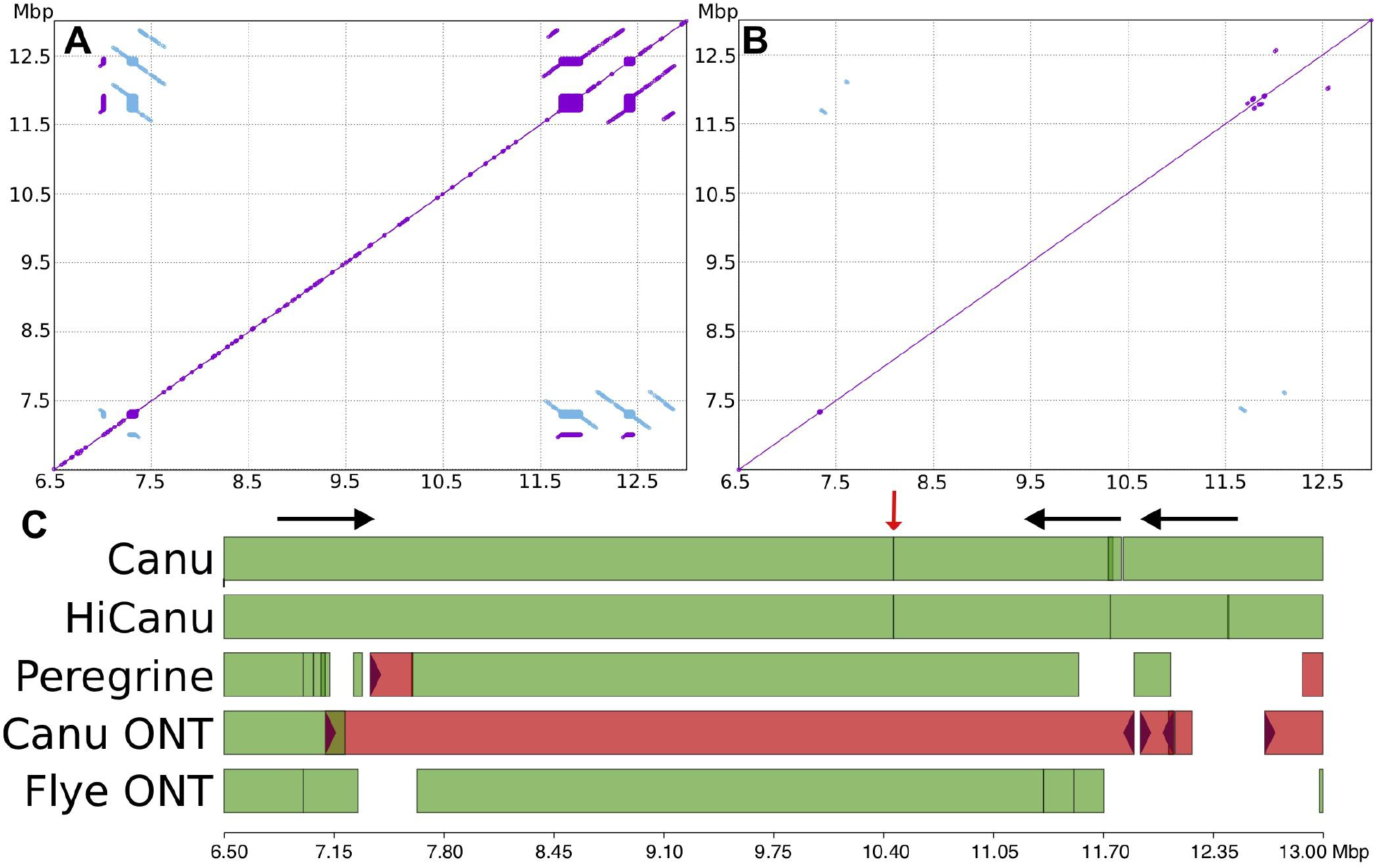
Chr8 β-defensin cluster repeat structure and assembly comparison. Top: Nucmer self-alignment dot plots (Kurtz et al. 2004) of the CHM13 reference defensin region at different alignment stringencies (Methods). A: greater than 7 kbp repeats at 98% identity. B: greater than 7 kbp repeats at 99.9% identity. Purple/blue indicates same/reverse strand matches. C: Icarus (Mikheenko et al. 2016) visualization of contig alignments from both HiFi-based (Canu, HiCanu, Peregrine) and ultra-long Nanopore-based assemblies (Canu ONT and Flye ONT (Kolmogorov et al. 2019)) produced by QUAST (Gurevich et al. 2013). White space in the alignment figure indicates the assembly was fragmented into short contigs (<50 kbp). Red color indicates mis-assembled contigs. The HiCanu assembly breaks at two of three segmental duplication instances which share high sequence similarity (black arrows) and at a region of systematic HiFi coverage depletion (red arrow).

The rightmost contig breaks in the HiFi assemblies are likely due to the presence of long, nearly identical repeats, which would require either longer reads or a careful examination of repeat copy number to resolve. We also investigated the fragmentation of HiCanu and Canu contigs at position 10.4 Mbp, which is not part of any observed repeat structure. Alignment of the raw HiFi reads onto this region with Minimap2 (Li 2018) revealed the presence of a 450 bp region covered by only two HiFi reads (Supplementary Figure 12), with a coverage drop present in both the 10 and 20 kbp HiFi libraries. This coverage drop is flanked by a >250 bp simple-sequence repeat (AAAGG). Suspecting a possible bias in the HiFi datatype, we further examined chromosome X, for which we have a complete CHM13 reference sequence available (Miga et al. 2019). On this chromosome, we identified at least four additional cases of HiFi coverage dropout, with all four instances associated with long, low-complexity (A/G or T/C-rich) sequences. As our HiFi assembly of chromosome X is split into just 13 large contigs, this coverage bias appears to be a current weakness of the HiFi chemistry.

## Discussion

We have demonstrated that HiCanu is capable of generating the most accurate and complete human genome assemblies to date, and our approach to mitigating sequencing errors in HiFi reads is general enough to be applied to other applications, such as metagenomic assembly. HiFi data excels in resolving large highly-similar (but non-identical) repeat instances. The remaining unresolved sequences seem to primarily represent satellite repeats (Supplementary Figure 6). In particular, Figure 2 illustrates that HiCanu’s reconstruction of human chromosomes 1, 7, 9 and 16 notably improves continuity over the previous assembly of ultra-long Nanopore reads (Koren et al. 2017; Kolmogorov et al. 2019; Shafin et al. 2019). These chromosomes contain several segmental duplications exceeding 200 kbp in length, requiring high-fidelity reads to identify variants and separate the individual copies. HiFi data further enabled draft assemblies of nine centromeric regions, which are one of the final challenges of automated telomere-to-telomere assembly. Assembly of other centromeric regions is likely limited by a low frequency of unique markers as compared to current HiFi read lengths. In contrast, chromosome X has the highest count of large (>20 kbp) near-identical (>99.9%) repeats (Bailey et al. 2002) that were better resolved by long, spanning Nanopore reads. Thus, the two technologies are complementary at present, and the best technology depends on the specific characteristics of the repeats and haplotypes being assembled.

When choosing HiFi, the library size should also be considered when beginning a sequencing project. Since HiFi read accuracy depends on the size of the sequenced fragments (shorter equals more passes and higher accuracy), one should consider the relative importance of read length versus accuracy. A metagenomic project may aim for shorter, higher accuracy reads to confidently identify low-abundance strains, while a vertebrate genome project may benefit from longer reads to span mid-sized identical repeats. We also identified an apparent bias in the current HiFi chemistry at low-complexity A/G (T/C) repeats, leading to coverage drops and assembly fragmentation. This issue warrants further investigation and may limit the applicability of HiFi sequencing to genomes with large stretches of such repeats. Thus, identifying optimal sequencing strategies and developing methods that can combine multiple technologies remains an area for future research.

HiCanu’s approach to read correction permits considering only the highest identity overlaps during contig construction. While HiCanu is not as fast as some of the competing methods, we note that the number of CPU hours required for assembly of a human genome is under 4,000, which could be completed on any modern cloud platform in less than a day for a few hundred dollars. This is 30-fold less than recent Oxford Nanopore assemblies that required more than 100,000 CPU h (Jain et al. 2018b; Shafin et al. 2019). At the time of writing, the most computationally expensive step of HiFi analysis is generating the data itself, since each individual HiFi read represents a consensus of multiple, aligned sequences of the same DNA molecule. Coupled with the instrument runtime and sequencing cost, HiCanu is a small fraction of the total project cost and duration (Supplementary Note 7).

While HiCanu’s diploid assemblies include long and accurate haplotype blocks of very high quality (QV50), they still represent pseudo-haplotypes (i.e. they do not preserve phase across their entire length). Canu also does not assign contigs to haplotypes, and requires postprocessing with a tool such as Purge_dups (Guan et al. 2020) to split the diploid assembly into primary and alternate alleles. While recent studies have successfully integrated HiFi data with additional long-range linkage information (Garg et al. 2019; Porubsky et al. 2019), we do not expect that significant improvements in phasing can be achieved by HiFi-only assemblies without an increase of HiFi read lengths. One option is post-processing of HiCanu assemblies by a haplotype-aware scaffolder, such as FALCON-Phase (Kronenberg et al. 2019), which could potentially correct haplotype switch events and deliver further improvements to phasing accuracy and assembly continuity. In general, we feel that HiFi contigs combined with Hi-C phasing and scaffolding is a promising recipe for phased telomere-to-telomere vertebrate genome assembly, and we plan to integrate these data types in future versions of Canu.

## Methods

### Mitigating errors in HiFi data

While HiFi reads are highly accurate compared to other long-read sequencing technologies, they are not error-free, which complicates the identification of reads originating from the same genomic loci during assembly. To identify and remove false read overlaps, we sought to increase the accuracy of HiFi data via read correction.

Wenger *et al*. observed that the majority of HiFi errors are in homopolymers, where the number of individually repeating nucleotides is incorrectly counted (Wenger et al. 2019). To lessen the impact of such errors, HiCanu modifies the input reads by compressing every homopolymer to a single nucleotide. Our approach is similar to run-length encoding (RLE), which has been previously applied to 454 (Miller et al. 2008), Pacbio CLR (Li 2016; Ruan and Li 2020; Li 2018) and Oxford Nanopore (Shafin et al. 2019) reads. However, HiCanu does not explicitly store the lengths of the compressed homopolymer stretches, and instead reverts back to the uncompressed reads when needed.

Although the transition to homopolymer-compressed sequence space can reduce the specificity of the read alignment search, the corresponding reduction in the number of observed errors in the reads allow for a more restrictive alignment identity threshold (based on empirical analyses we require a minimum overlap identity of 99%). Subsequent steps are performed on the homopolymer-compressed sequences, while the detailed correspondence between positions of original and compressed versions is generated on the fly when necessary. Compressed reads are first subjected to Overlap-Based Trimming (Koren et al. 2017). While this step does not affect the majority of HiFi assemblies, we enable it by default as a precautionary measure since it considerably improved assembly contiguity on one recent dataset (HG002 Sequel II System with pre-2.0 Early Access Chemistry 15 kbp library available from https://github.com/human-pangenomics/HG002_Data_Freeze_v1.0). This improvement suggests that a significant fraction of reads were structurally incorrect due to a low-quality sequencing library. Since other libraries did not show this problem, it is likely future versions of the HiCanu pipeline can skip this step and reduce runtime by more than 60%.

To further reduce the influence of the errors in compressed HiFi reads we have updated the Overlap Error Adjustment (OEA) module of Canu (Holt et al. 2002; Koren et al. 2017). This module identifies errors in individual reads by jointly considering all of their overlapping reads. Every such read votes for the nucleotides at the positions that it covers based on the pairwise alignment of the overlapping regions. A read’s position is considered erroneous if no other reads support the original sequence and the majority of votes agree on a particular change (by default more than 50% and at least 7 if there is a read supporting the original sequence). After the corrections are introduced the alignment scores of the overlaps are recomputed, but the actual read sequences stored within the assembler are not modified as doing so would invalidate the previously computed overlap coordinates. Although our naive approach may not always be able to correct errors within highly-similar genomic repeats, such events are rare due to the low number of errors in compressed HiFi reads and the high identity threshold used for gathering candidate overlaps.

Manual investigation of read alignments during HiCanu development revealed a previously unreported error mode in HiFi reads: incorrect repeat unit counts within microsatellite repeat arrays. Since the incorrect repeat counts are systematic and often supported by multiple reads, the conservative strategy described above is not able to correct them. Recognizing this, we modified the OEA procedure for recomputing overlap alignment scores to ignore sequence differences flanked by a microsatellite repeat in either read. Namely, the difference is ignored if 5 out of 6 non-overlapping flanking k-mers are the same for any k ranging between 2 and 6 on either side (starting at 0 to k-1 bp from the difference). We note that this phenomenon deserves a deeper investigation and our strategy can be improved to capture additional genomic differences, which are ignored by the current approach.

We evaluated the contribution of each of the above corrections using the recently completed CHM13 chromosome X (Miga et al. 2019) as a reference. Raw, compressed, corrected, and masked 20 kbp HiFi reads were mapped and the mappings filtered to retain high-confidence alignments (Supplementary Note 1). Figure 1 shows the resulting alignment identity values, with each correction step boosting the identity of the aligned sequence. Each step (compression, correction, masking) contributes to this improvement (Supplementary Table 1), and the combined effect is most dramatic for the number of perfect alignments. While almost no (<1%) raw HiFi reads map error-free, 97.23% of the compressed, corrected, and masked reads map without a single difference. Because we did not control for reads mapping from other chromosomes, and the chromosome X sequence itself is not error free, this likely represents a lower bound on the percentage of error-free reads.

### Bogart modifications

The Bogart module constructs a set of draft contigs from read overlap information. A detailed description is given in (Koren et al. 2017). We describe here the modifications made for HiFi data.

#### Overlap identity threshold

Canu’s initial overlap search uses a relaxed identity threshold to account for varying error rates between samples. Since overlap identities are changed by OEA, and to avoid considering falsepositive overlaps, Bogart first attempts to select a higher overlap identity threshold. Previously, Canu computed the identity of the best scoring overlap on each side of every read (where score is defined as the number of matching bases in the overlap alignment) and set a threshold based on the median and MAD of the computed values (Koren et al. 2017). However, during the development we realized that this way of computing the threshold was not informative for highly accurate reads because both the median and MAD were 100% across all tested datasets. Additionally, with the number of matching bases as a score, the read delivering the highest scoring overlap could come from a different haplotype in genomes with low heterozygosity. As a result, the selected threshold could inadvertently reflect the heterozygosity level of the organism rather than the accuracy of the reads. Based on empirical testing, we opted for an alternative two-step procedure. First, all overlaps with identity below a fixed value T (default 99.97% or 3 differences in 10,000bp) are dropped. This step is aimed at removing from consideration the majority of the cross-haplotype overlaps even for low-heterozygosity organisms, e.g. human heterozygosity rate of 0.1% (1000 Genomes Project Consortium et al. 2012). Next, the identities of the highest scoring overlaps are collected as before and the final threshold is set as the 90th% percentile of this sample. It is possible that 99.97% is too stringent given higher error reads. We could detect this condition when the 90th percentile is too close to 99.97% and re-run the overlap filtering. However, on all datasets evaluated to date, the chosen identity threshold was 100%. To support the desired overlap filtering stringency, the Canu codebase had to be modified to increase the precision with which the overlap identity values are stored.

#### Handling heterozygous differences

Bogart uses the filtered overlaps to identify and eliminate the reads likely representing sequencing artifacts and then constructs the *best overlap graph* (Miller et al. 2008), using the same overlap scoring function as before. This graph consists of the best scoring overlap off both the 5’ and 3’ ends of each read, and the non-branching paths within this graph form the preliminary layouts (arrangements of reads) that we refer to as *greedy contigs*. Bogart then inspects each greedy contig for long repeat instances that could have been incorrectly traversed. Repeats are detected by considering overlaps between the reads within and outside of the contig. If a suspected repeat has no reads spanning it, or there is a similar-length alternate read overlap, it is broken at the repeat boundary to avoid potential assembly errors as in (Koren et al. 2017) to form the final *draft contigs*.

HiCanu aims to reconstruct long pseudo-haplotype contigs (Vinson et al. 2005; Chin et al. 2016)—potentially switching between paternal and maternal alleles—and capture the alternative regions as shorter contigs. Unfortunately, the original Bogart approach described above led to the classification of extended homozygous regions within greedy contigs as unspanned repeats, leading to fragmentation of the pseudo-haplotypes (Supplementary Figure 13). In Canu, this behavior had been affecting only genomes with >1% heterozygosity, since below this threshold most heterozygous differences were implicitly hidden by the relatively permissive threshold on overlap identity. With the high accuracy HiFi data, and correspondingly high overlap identity threshold, this over-fragmentation became an issue even for a human levels of heterozygosity.

In HiCanu, Bogart has an additional step to identify contigs representing alternative alleles within the set of greedy contigs, which we refer to as *bubble contigs*. As suggested by the name, the bubble contigs are related to the bubble subgraphs, typically considered by most assemblers. Candidate bubbles are found by identifying reads in each contig that have overlaps to some other, larger, contig. A read within a smaller contig can be placed in the larger contig if the overlaps between it and the reads in the larger contig are below a specified threshold of similar quality to the previously incorporated overlaps (0.1% by default). If the placements for both the first and last reads of a candidate contig are correctly oriented and placed at approximately the correct distance in the larger contig (75–125% of the candidate contig size), the candidate contig is flagged as a bubble and its reads are excluded from later repeat detection. This avoids fragmentation of otherwise structurally correct pseudo-haplotype contigs. Similar strategies have previously been used in short-read assembly (Pevzner et al. 2001; Zerbino and Birney 2008; Li et al. 2010; Gnerre et al. 2011), scaffolding metagenomes (Koren et al. 2011; Ghurye et al. 2019) and long-read assembly (Chin et al. 2016). Bubble contigs are also explicitly marked in the final output; however, because placements are not always found, especially for longer, more heterozygous alleles, we recommend using a postprocessing tool such as Purge_dups (Guan et al. 2020) to classify alternate alleles and remove any false duplications.

### Consensus calculation

A consensus sequence is computed for all contigs using the uncompressed reads (trimmed to their good regions identified in compressed space). Canu originally used the layout produced by Bogart to estimate the position of each read within the contig and align it only to that location. Since the read layouts are now in homopolymer-compressed space, this strategy is unable to locate the read in uncompressed space. Instead, we compute the correspondence of each position in the compressed read to the original. This is used to update the read positions within the contig and expand the layout to uncompressed space. A modified version of the PBdagcon algorithm (Chin et al. 2013), with improved support for long contig sequences, is used to compute the final consensus sequence. Further consensus gains are likely possible using a more sophisticated approach for predicting homopolymer run length, similar to MarginPolish (Shafin et al. 2019).

### Software commands

HiCanu was run using Canu branch hicanu_rc with the following commands:

~~~
       canu -assemble -p asm -d asm genomeSize=G -pacbio-hifi reads.fastq.gz
~~~

with G=3.1g for human and 150m for *D. melanogaster*. This required 131 CPU h and 16 GB of memory for *D. melanogaster*, 2780 CPU h and 66 GB of memory for CHM13 10 kbp library, 5000 CPU h and 119 GB of memory for CHM13 20 kbp, 3999 CPU h and 62 GB of memory for HG002, and 5,233 CPU h and 50 GB of memory for HG00733.

For the standard Canu assembles, Canu branch hicanu_rc ran with the following command:

~~~
       canu -p asm -d asm genomeSize=G correctedErrorRate=0.015 batOptions=“-eg 0.01 -eM 0.01 -dg 6 -db 6
       -dr 1 -ca 50 -cp 5” -pacbio-corrected reads.fastq.gz
~~~

with G=3.1g for human and 150m for *D. melanogaster*. This required 232 CPU h and 12 GB of memory for *D. melanogaster*, 3,524 CPU h and 80 GB of memory for CHM13 20 kbp library, and 3,836 CPU h and 47 GB of memory for HG00733.

For CLR data Canu branch hicanu_rc was run with the following command:

~~~
       canu -p asm -d asm genomeSize=150m corOutCoverage=100 batOptions=“-dg 6 -db 6 -dr 1 -ca 500 -cp 50” -pacbio-raw reads.fastq.gz
~~~

All HiFi assemblies required less than 12 wall-clock hours on the NIH Biowulf cluster quick partition with all jobs using <120 GB RAM. We estimated the cost of an AWS run using the c5d.18xlarge instance which costs $3.456/hr. Assuming 4 reserved nodes and an average runtime of 4200 CPU h, the run would complete in 14.5 hours. We increase this by a factor of 1.5 to account for any non-parallelized steps at a cost of $3.456 * 4 * 22 = $304. This cost could be reduced further if additional nodes were spun up on-demand for the parallel portions of compute and spun down when not needed. We omit this from the estimate for simplicity.

Peregrine Assembler & SHIMMER ASMKit (0.1.5.3) was run with the command:

~~~
yes yes | python3 /data/korens/devel/Peregrine/bin/pg_run.py asm \
    chm13.list 24 24 24 24 24 24 24 24 24 \
    --with-consensus --shimmer-r 3 --best_n_ovlp 8 \
    --output ./
~~~

This required 7 CPU h and 29 GB of memory for *D. melanogaster*, 32 CPU h and 347 GB of memory for CHM13 10 kbp library, 58 CPU h and 449 GB of memory for CHM13 20 kbp library, 55 CPU h and 407 GB for HG0002, and 63 CPU h and 477 GB for HG00733.

#### Commands for β-defensin validation

MUmmer 3.23 was used to identify repeats with the command:

~~~
       nucmer --maxmatch --nosimplify
       delta-filter -i 98 -l 10000
~~~

And high-stringency repeats:

~~~
       nucmer --maxmatch --noextend --nosimplify -l 500 -c 1000
       delta-filter -i 99.9 -l 10000
~~~

QUAST alignments were generated as:

~~~
       Quast.py --large --skip-unaligned-mis-contigs
~~~

#### Commands for RepeatMasker

RepeatMasker version 4.1.0 was run with the commands:

~~~
       RepeatMasker -pa 8 -q -species=mammal -xm -dir=asm.out asm.fasta
~~~

on each contig ≥50 kbp in the assembly. Centromeric arrays were identified by taking all hits marked as Satellite/centr and merging any hits within 100 bp of each other using bedtools (Quinlan and Hall 2010). Resulting arrays greater than 800 kbp were reported. There were nine internal arrays whose start and end coordinates were at least 500 kbp away from a contig end. These initial coordinates were manually curated based on reference alignments and are reported in (Supplementary Table 10).

#### Commands for MHC typing

HLA*LA version commit 24930adadb0d2b6bcd69a271401dfc88a5d09f4d was run with the commands:

~~~
       HLA-ASM.pl --use_minimap2 1 --assembly_fasta $asm --sampleID $prefix --workingDir ‘pwd’/$prefix --truth reference_HLA_ASM/$truth
~~~

where $asm was the assembly, $prefix was a unique identifier, and $truth was either truth_HG002.txt or truth_HG00733.txt.

#### Commands for Purge_dups

Purge_dups version commit 8f580b41e6aa20c99383d6ff19b8689e93d7490e was run with the commands:

~~~
       python pd_config.py asm.fasta ‘pwd’ <pb folder> <10x folder left blank> asm
       minimap2 -I6G -xasm5 -DP asm.split asm.split > asm.split.self.paf
       minimap2 -I6G -xmap-pb asm.fasta $line > pb.$jobid.paf (for each HiFi cell)
       pbcstat pb.*.paf
       calcuts PB.stat > cutoffs 2>calcults.log
       purge_dups -2 -T cutoffs -c PB.base.cov asm.split.self.paf > dups.bed 2> purge_dups.log
       get_seqs dups.bed asm.fasta > purged.fa 2> hap.fa
~~~

For *D. melanogaster*, an incorrect threshold was computed for the cutoffs due to the entire genome being separated and so the cutoffs were manually adjusted to be

~~~
       50 1 1 115 2 200
~~~

The purged.fa output was then used as the primary set reported in the tables. To obtain the alternate set, we ran a second round of Purge_dups using hap.fa as the input assembly instead.

#### Commands for Merqury

Merqury version commit 154610d19ee6f4fead77da077af1ed7abdbe8866 was used. For each assembly and read set, canonical k-mers were built using meryl available as a binary within Canu:

~~~
       meryl count k=<k-size> <reads.fastq.gz> output <genome>.k<k-size>.meryl
       meryl count k=<k-size> <asm.fasta> output <asm>.k<k-size>.meryl
~~~

using k=21 for humans and k=18 for *D. melanogaster* based on (Fofanov et al. 2004). QV and k-mer completeness were obtained with:

~~~
       eval/spectra_cn.sh
~~~

which converts k-mer Jaccard to distance as in (Ondov et al. 2019) and to a Phred score (Ewing and Green 1998). Haplotype blocks were estimated by first building parent-specific k-mer databases. K-mers in each parental dataset were counted as above, then subtracted to obtain parent-specific k-mers, and finally intersected with the child (in the case of human datasets where child Illumina data was available) with:

~~~
       trio/hapmers.sh
       trio/phased_block.sh
~~~

For further information see Supplementary Note 2 and https://github.com/marbl/merqury/wiki..

#### Commands used for QUAST

QUAST 5.0.2 ran with the command:

~~~
       quast.py <asm> -o quast_results/<asm> -r <reference> -t 16 -s --large
~~~

Variants were filtered using the pipeline from (Shafin et al. 2019) to filter errors in varying sites, including known SVs (HG002 only available from GIAB (Zook et al. 2019) at ftp://ftptrace.ncbi.nlm.nih.gov/giab/ftp/data/AshkenazimTrio/analysis/NIST_SVs_Integration_v0.6/HG002_SVs_Tier1plusTier2_v0.6.1.bed):

~~~
python3 reference/quast_sv_extractor.py -q quast_results/<asm>/contigs_reports/all_alignments*tsv -c reference/centromere.bed -d reference/GRCh38_marked_regions.bed -s reference/empty
~~~

and

~~~
       python3 reference/quast_sv_extractor.py -q quast_results/<asm>/contigs_reports/all_alignments*tsv -c reference/centromere.bed -d reference/GRCh38_marked_regions.bed -s reference/HG002_SVs_Tier1plusTier2_v0.6.1.bed
~~~

for HG002. We used https://www.ncbi.nlm.nih.gov/assembly/GCF_000001215.4, filtered to remove any unassigned sequences for *D. melanogaster* (chr2L, chr2R, chr3L, chr3R, chr4, chrM, chrX, chrY only) and https://hgdownload.soe.ucsc.edu/goldenPath/hg38/bigZips/hg38.fa.gz filtered to exclude alts and unaligned sequences (chromosomes 1-22, X, Y, and MT only). Since no filtering file was available for *D. melanogaster*, we modified QUAST parameters to try to avoid false-positive misassembly counts with the command:

~~~
       quast.py <asm> -o quast_results/<asm> -r <reference> --large --min-alignment 20000 --extensive-mis-size 500000 --min-identity 90
~~~

#### Commands for BAC validation

We used the BAC validation pipeline available at https://github.com/skoren/bacValidation run with default parameters. This pipeline aligns reads using minimap2 (Li 2018) and parses the SAM (Li et al. 2009) format to generate summary statistics. Output BAC identity was computed as the median across all BACs marked as correctly resolved. BAC libraries were downloaded from NCBI (CHM13: https://www.ncbi.nlm.nih.gov/nuccore/?term=VMRC59+and=complete, HG00733: https://www.ncbi.nlm.nih.gov/nuccore/?term=VMRC62+and+complete). HiFi read alignments to the assembly and BAC sequences were visualized with IGV (Robinson et al. 2011)

#### Commands for identifying contig ends

Alignments were made between assemblies and GRCh38 using the following Minimap2 command:

~~~
       minimap2 --secondary=no -a --eqx -Y -x asm20 -s 200000 -z 10000,50 -r 50000 --end-bonus=100 -O 5,56 -E 4,1 -B 5
~~~

Contig ends that intersected segmental duplications were identified by parsing the CIGAR string to find the location of contig ends and then intersecting these locations with annotated segmental duplications plus 10kbp on either side from the UCSC genome browser using the following commands:

~~~
       bedtools slop -i {segdups.bed} -b 10000 | bedtools merge -i - > {expanded.segdups.bed} && bedtools
intersect -a {contig.ends.bed} -b {expanded.segdups.bed}
~~~

## Data Availability

We have posted the downsampled datasets, generated assemblies, and corrected CHM13 BAC sequences at https://obj.umiacs.umd.edu/marbl_publications/hicanu/index.html. When available, previously published assemblies were downloaded and used. This included Oxford Nanopore UL Canu assemblies presented by (Shafin et al. 2019) for HG0002 (80x Guppy HAC 2.3.5) and HG00733 (50x Guppy HAC 2.3.5); Canu + Racon assembly presented by (Vollger et al. 2020); HG002 Canu assembly of HiFi reads presented by (Wenger et al. 2019); Oxford Nanopore Canu assembly for CHM13 (40x + 80x UL Guppy HAC 3.1.5) presented by (Miga et al. 2019); HiFi + Hi-C assemblies for HG002 presented by (Garg et al. 2019); HiFi + Strand-seq assemblies for HG0733 presented by (Porubsky et al. 2019). In the remaining cases, assemblies were run locally on the NIH Biowulf cluster.

The *D. melanogaster* HiFi data is available from NCBI at PRJNA573706 (SRR10238607, median:, 24.4 kbp mean: 24.4 kbp) and CLR (SRR9969843, median 13.3 kbp, mean 17.2 kbp). Due to the high coverage, this dataset was downsampled to 40× HiFi data and 200× CLR data. These coverages represent approximately 25% of the full run output. Since the exact parents of the F1 were not available, we used the previously generated short-read sequencing for binning and analysis (A4: SAMN00849823, ISO1: SRR6702604). The CHM13 Nanopore data is available at https://s3.amazonaws.com/nanopore-human-wgs/chm13/nanopore/rel3/rel3.fastq.gz, HiFi reads in PRJNA530776 (10 kbp: SRR9087597-SRR9087600; 20 kbp: SRR11292120-SRR11292123), and Illumina at https://github.com/nanopore-wgs-consortium/CHM13#10x-genomics-data. The HG002 Nanopore data is available at https://s3-us-west-2.amazonaws.com/human-pangenomics/index.html, HiFi at SRX5327410. HG002 and parent Illumina data is available from GIAB (Zook et al. 2016) at https://github.com/genome-in-a-bottle/giab_data_indexes, we only used the 2×250 datasets. The HG00733 Nanopore data is available at https://s3-us-west-2.amazonaws.com/human-pangenomics/index.html, HiFi at ERX3831682. Illumina data for HG00733 and parents were downloaded from the 1000 Genome consortium at https://www.internationalgenome.org/data-portal/sample (1000 Genomes Project Consortium et al. 2012).

The CHM13 chromosome 8 reference assembly is available at https://github.com/nanopore-wgs-consortium/CHM13#downloads.

## Acknowledgements

This work was supported, in part, by the Intramural Research Program of the National Human Genome Research Institute, National Institutes of Health (SN, BPW, AR, AMP, and SK) as well as grants from the U.S. National Institutes of Health (NIH HG010169 and HG002385 to EEE), National Human Genome Research Institute (NHGRI R21 1R21HG010548-01 to KHM), and National Institute of General Medical Sciences (NIGMS 1F32GM134558 to GAL). EEE is an investigator of the Howard Hughes Medical Institute. This work utilized the computational resources of the NIH HPC Biowulf cluster (https://hpc.nih.gov). We thank Pacific Biosciences for providing the *D. melanogaster* F1 andCHM13 20 kbp data.

## Disclosure Declaration

RG is an employee and shareholder of Pacific Biosciences. EEE is on the scientific advisory board of DNAnexus, Inc. All other authors have no competing interests to declare.

## Notes

#### Summary of Updates

This version fixes the accession for CHM13 HiFi data

https://obj.umiacs.umd.edu/marbl_publications/hicanu/index.html

