## Supplementary Notes, Figures, Tables for "HiCanu: accurate assembly of segmental duplications, satellites, and allelic variants from high-fidelity long reads"

- Supplementary Note 1: Evaluation of HiFi read accuracy 2
- Supplementary Note 2: Using k-mers for assembly evaluation and identifying haplotype-blocks within diploid assemblies 3
- Supplementary Note 3: CHM13 Challenge BAC Validation 4
- Supplementary Note 4: Identification of low-coverage gaps 5
- Supplementary Note 5: Human repeat modeling 6
- Supplementary Note 6: Centromere analysis and validation 7
- Supplementary Note 7: Estimate of AWS costs 8
- Supplementary Figures 9
- Supplementary Tables 22

### Supplementary Note 1: Evaluation of HiFi read accuracy

We took all 20 kbp HiFi sequences from the CHM13 cell line and aligned them to the v0.7 chrX assembly. The reads were aligned with minimap2 (Li 2018) v2.17 with the command:

```
minimap2 -a -H -t 15 -x asm20
```

Reads were then homopolymer-compressed (and trimmed by Canu, though untrimmed reads showed a similar alignment identities) and aligned to the same reference. Finally, the compressed and trimmed reads were corrected by Canu's OEA module and aligned. Only primary unambiguous alignments were retained using the command:

```
samtools view -F 2304 -q 60
```

Then alignment identity values were computed by a custom script (https://github.com/snurk/bacValidation/blob/master/samToErrorRate.C), optionally ignoring the differences flanked by microsatellite repeat arrays. Only reads with a single alignment covering at least 99% of their length with identity >98% were evaluated. Note that identity may vary across the genome and these are summary statistics, as both sequencing errors and the correction procedure are region-specific.

### Supplementary Note 2: Using k-mers for assembly evaluation and identifying haplotype-blocks within diploid assemblies

We used Merqury (Rhie et al 2020) to estimate assembly k-mer level QV, completeness, and phase block statistics. We computed an optimal k-mer size of 18 for the *D. melanogaster* and 21 for human, as in Fofanov et al., with a collision rate of 0.001. This calculation is automated in Merqury's `best_k.sh script.`

**QV and k-mer completeness**

The assembly distance from the read set is defined as the number of k-mers found only in the assembly divided by the total number of k-mers found in the assembly. This can be converted to a distance as in Ondov et al. 2019:

$$P = (K_{shared} / K_{total})^{1/k}$$

and to a Phred scale QV as:

$$QV = -10 \log_{10} 1\text{-}distance$$

Completeness is measured as the fraction of reliable k-mers in the read set present in an assembly. To determine reliable k-mers, we build the histogram of k-mer counts (number of times a k-mer is seen in the read set (multiplicity) versus number of k-mers with a given multiplicity). The minimum reliable k-mer threshold is set to the lowest multiplicity with a positive slope in the histogram. The filtering is automated in Merqury, with `build/filt.sh.`

**Phase blocks**

Haplotype-specific markers were determined using parental specific k-mers as previously done for TrioBinning (Koren et al. 2018). K-mer databases were built for each parental read and subtracted to obtain parent-specific markers. Reliable k-mers for each parent-specific database are selected as above. These databases were used for the *D. melanogaster*. For human datasets, the parent-specific k-mer databases were further intersected with the child's F1 Illumina data to only include parent-specific markers which were inherited by the child.

Phase blocks were obtained by querying each k-mer found in the assembly to each parental k-mer database. A phase block is defined as having at least 2 haplotype-specific markers from the same haplotype, allowing short-range "switches" to the other haplotype. We defined a short-range switch as at most 100 consecutive markers within a 20 kbp region. When more than 100 markers from the other haplotype are found or they span more than 20 kbp, a new block is created. The new block starts at the first inconsistent marker found. The switch error rate is defined as the fraction of markers from the wrong haplotype within all phase blocks.

### Supplementary Note 3: CHM13 Challenge BAC Validation

We inspected all 15 of the CHM13 BACs which were not correctly resolved by the HiCanu assembly. We mapped all 20 kbp HiFi reads to the BACs and to the HiCanu assembly with minimap2:

```
minimap2 -t 32 -ax map-pb -r 2000 -m 3000 <reference.fasta> *.fastq.gz
```

We also checked the BACs for sequence similarity to the vector and *E. coli* sequences. We concluded that 11 of the sequences were incorrect or highly suspicious:

- AC278245.1 Low complexity sequence likely expanded or contracted within the BAC. (likely error)
- AC278709.1 BAC includes 2047 bp of cloning vector (error).
- AC278859.1 BAC includes 2047 bp of cloning vector (error).
- AC278792.1 Soft clipped 331bp (left) of sequence in the BAC that matches many other BACs in BLAST at perfect identity, untrimmed vector (error), and soft clipped (right) 4779 of cloning vector (error).
- AC278658.1 BAC includes 2047 bp of cloning vector (error).
- AC278258.1 BAC includes 872 bp of cloning vector (error).
- AC278368.1 BAC includes 2047 bp of cloning vector (error).
- AC279108.1 Insertion of 246 bases in the assembly supported by all HiFi data (likely error), Supplementary Figure 2
- AC278968.1 Lots of base mismatches and a large SV being called by all HiFi data. (likely error), Supplementary Figure 3
- AC278708.1 Soft clipped 331bp of sequence in the BAC that matches many other BACs in BLAST at perfect identity, likely untrimmed sequence of some kind. Clipped identically in GRCh38.p13 and HiFi and UL assemblies (likely error), Supplementary Figure 4
- AC278985.1 Lots of base mismatches and a large amount of soft clipping (2045). Clipped identically in GRCh38.p13 and HiFi and UL assemblies (likely error), Supplementary Figure 5

The adjusted CHM13 BAC resolution stats can be found in Supplementary Table 5.
We claim that only four BAC regions were unresolved by the HiCanu assembly:

- AC279099.1 Soft-clipped BAC extends past contig end
- AC279073.1 Lots of mismatches across the whole BAC and 7345 bp of soft clipped sequence extends past the contig end.
- AC270468.1 BAC extends past the contig end.
- AC278922.1 BAC split across two contigs

### Supplementary Note 4: Identification of low-coverage gaps

All HiFi reads were mapped to the chr8 and chrX T2T assembly with minimap2 with the commands:

```
minimap2 -a -H -t 15 -x asm20
```

The HiCanu assembled contigs were also mapped to the T2T assembly with the same commands. We identified regions with <= 2-fold HiFi coverage coinciding with a break in the contig mapping. The regions were visually inspected in IGV to confirm simple-sequence repeats.

#### Supplementary Note 5: Human repeat modeling

We previously developed a model of human assembly continuity to predict the effects of read length and accuracy. Increased read length allows for more repeats to be spanned by reads that are uniquely anchored on either side, while increased read accuracy allows for more repeats to be separated based on sequence variants within the repeat. Using PacBio CLR or Nanopore data, existing algorithms have been able to separate repeats that are up to 98% identical at the sequence level by detecting variants within the repeats. Assuming a 20 kbp read length, this model predicts that improving repeat separation from 98% to 99.9% identical repeats would boost assembly NG50 by 1.8-fold (N such that 50% of the genome size is assembled in contigs of this size or greater). Thus, because of their improved ability to separate repeats, highly accurate 20 kbp reads are predicted to rival noisy 200 kbp reads in terms of assembly continuity (Supplementary Figure 7).

### Supplementary Note 6: Centromere analysis and validation

HiFi data was aligned to the HiCanu assembly with pbmm2 v1.1.0 and the command:

```
pbmm2 align --preset SUBREAD -N 50 --min-length 3000 -r 50000 {input.ref} {input.reads} | samtools view -u -F 2308 - | samtools sort -o {output}
```

ONT rel3 data from (Miga et al. 2019) was aligned to the HiCanu assembly with minimap2 with the command:

```
minimap2 -ax map-ont -k15 -w5 -N 50 -r 10000 <asm.fasta> <reads.fastq>
```

and filtered unique markers as in (Miga et al. 2019). The resulting coverage was visualized with IGV and repeats within each array annotated using RepeatMasker.

The 606 kbp D19Z1 α-satellite array within the chromosome 19 centromeric repeat is composed of a 2.25 kb (13-mer) HOR unit (Hulsebos et al. 1988), completing the repeat representation from the partial (10-mer) sequence available in Genbank (AJ295045.1). Further, we demonstrate that intra-array homogeneity is high, with units of the 13-mer HORs within the D19Z1 array 97.5% identical on average (Supplementary Figure 9a). The 3.96 Mbp D19Z3 α-satellite array is predicted to consist primarily of a dimeric HOR unit (Baldini et al. 1989). While the dimeric array on chromosome 1, 5, and 19 centromeres (pC1.8 clone, Genbank M26919 and M26920) has demonstrated similar hybridization patterns (Baldini et al. 1989; Finelli et al.1996), these sequences shared only 93% sequence identity to the HiCanu D19Z3 HOR array, on average. This supports previous findings that chromosome-specific sequences are present in D19Z3 to distinguish these arrays at the sequence level (Pironon, et al. 2010). Here, we report a new dimeric α-satellite sequence for D19Z3 that is 95.7% identical, on average, between HOR units in the array. (Supplementary Figure 9b). Finally, the ~300 kbp D19Z2 array consists of two different HORs, but these repeat structures had not been characterized before. However, they share sequence similarity with the GJ211883.1 (cen5_2) and GJ211884.1 (cen5_4) alpha satellite reference models in GRCh38. Using the HiCanu assembly and the underlying raw HiFi data, we identified two complex D19Z2 HOR structures that are, themselves, comprised of α-satellite HORs. These complex HOR structures are organized as follows: 7-, 8-, 6-, 7-, 4-mer, and 6-, 6-, 3-, 4-, 4-, 7-, 4-, 4-, 2-mer (Supplementary Figure 10).

### Supplementary Note 7: Estimate of AWS costs

Assuming a 30-hr movie (https://www.pacb.com/products-and-services/sequel-system/latest-system-release/), acquiring CHM13 data on four SMRTcells would require 120 hours. We obtained 3/6 NA12878 raw bam files for the subreads and converted them to HiFi using the CCS command v3.4.1:

```
        ccs --maxLength 21000 --minPasses 3 --numThreads 32 --polish --minPredictedAccuracy 0.99 input.bam
output.bam
```

The jobs took on average 7,136 CPU h and 13 GB memory. Thus, we estimated a total of 42,817 CPU h to convert all six cells to HiFi reads. This corresponds to 199 hours on a 36-core node per cell. Using a c5d.9xlarge (1.728/hr) instance to allow storing large BAM files locally, this would cost $2,055. Finally, we estimate the HiCanu runtime on AWS to be 22 hours versus 2 hours for Peregrine. The total runtime with HiCanu is 341 and with Peregrine is 321, a difference of 6%. For cost, we ignore reagent and consumable cost and focus only on computation. We estimated above the cost to generate HiFi reads on AWS as $2,055 and HiCanu as $300 vs $10 for Peregrine. The total cost with HiCanu is $2,355 and with Peregrine $2,065, a difference of 15%.

### Supplementary Figures

**Supplementary Figure 1. *D. melanogaster* ISO1xA4 genomescope.**

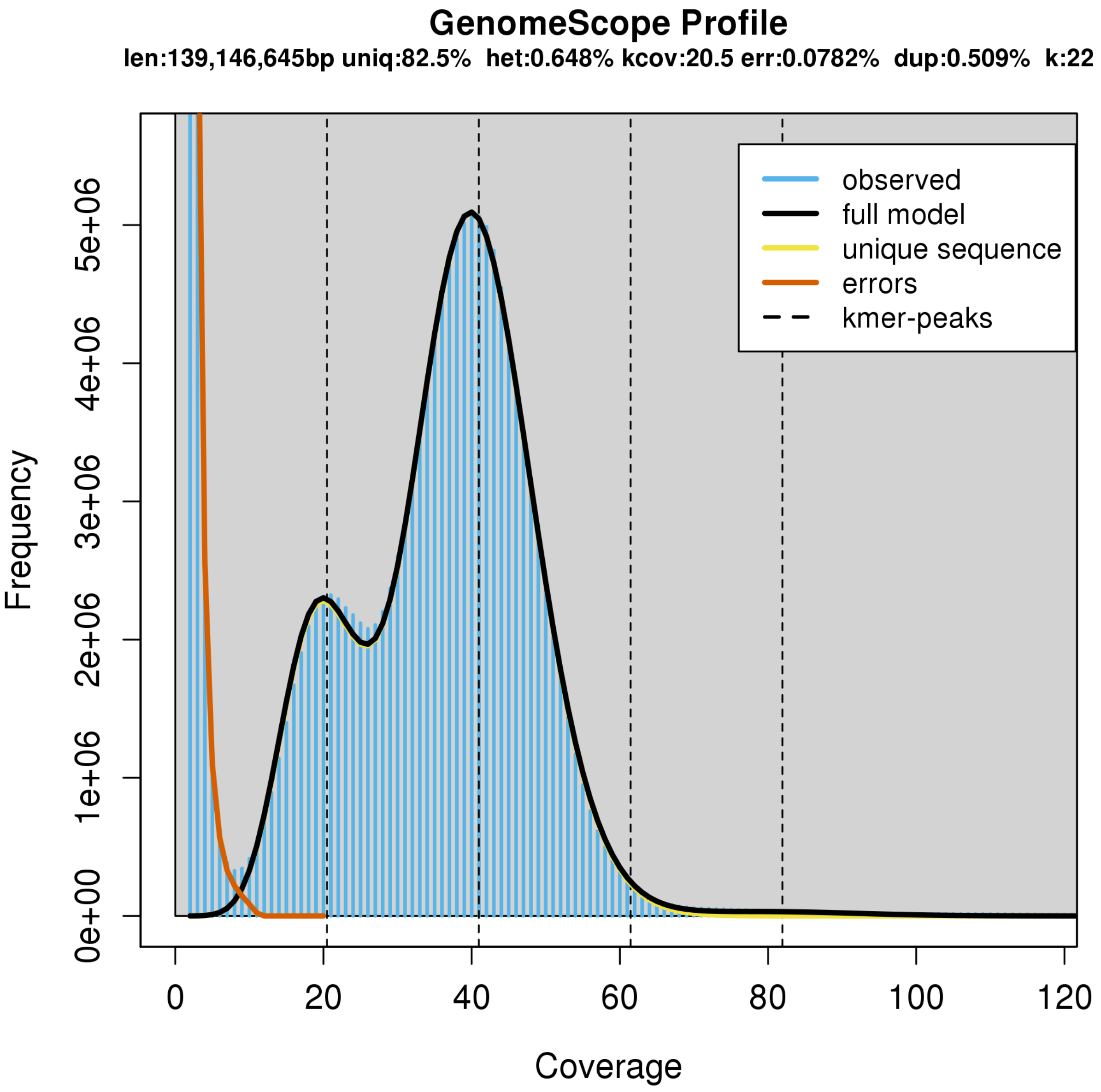


Estimate of genome size and heterozygosity using 22-mers found in the HiFi 24 kbp 40x downsampled library for *D. melanogaster* ISO1xA4

**Supplementary Figure 2. Evaluation of BAC AC279108.1.**

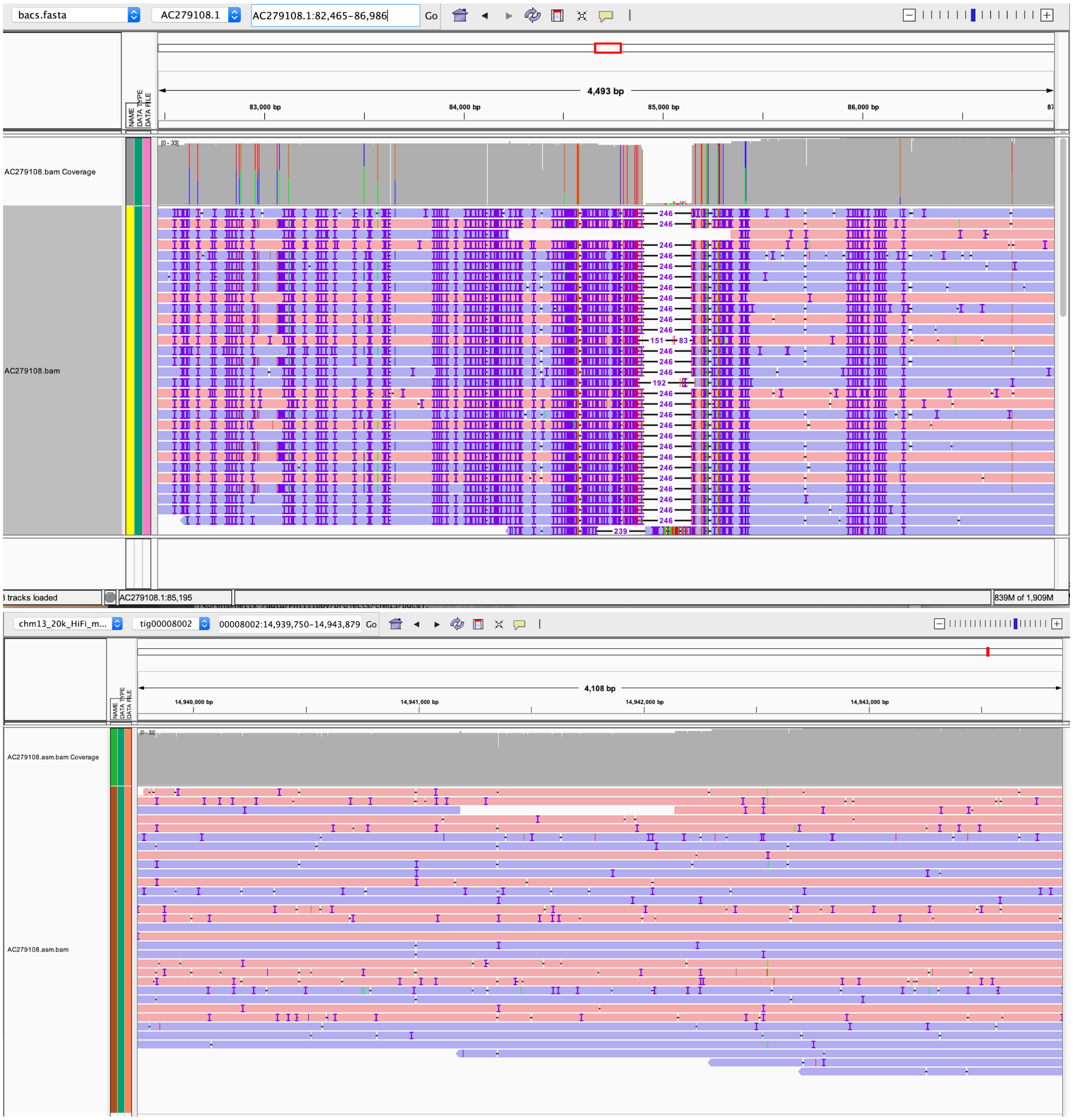


Top panel shows alignments of HiFi reads to the BAC sequence, bottom -- to the assembly. The top panel indicates a large SV called by the reads versus the BAC while the assembly is consistent with the data.

#### Supplementary Figure 3. Evaluation of BAC AC278968.1.

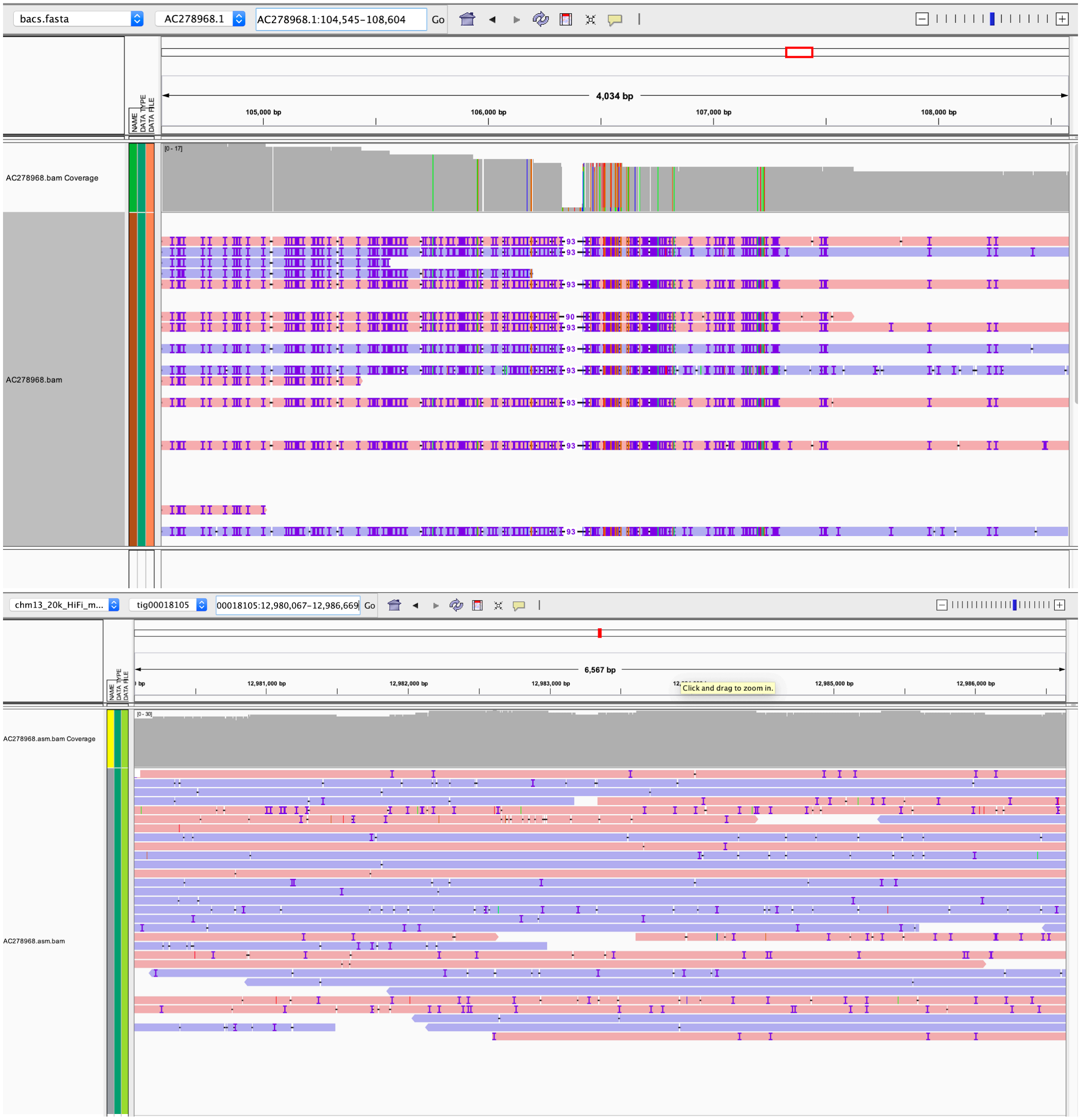


Top panel shows alignments of HiFi reads to the BAC sequence, bottom -- to the assembly. The top panel indicates a large SV called by the reads versus the BAC while the assembly is consistent with the data.

**Supplementary Figure 4. Evaluation of BAC AC278708.1.**

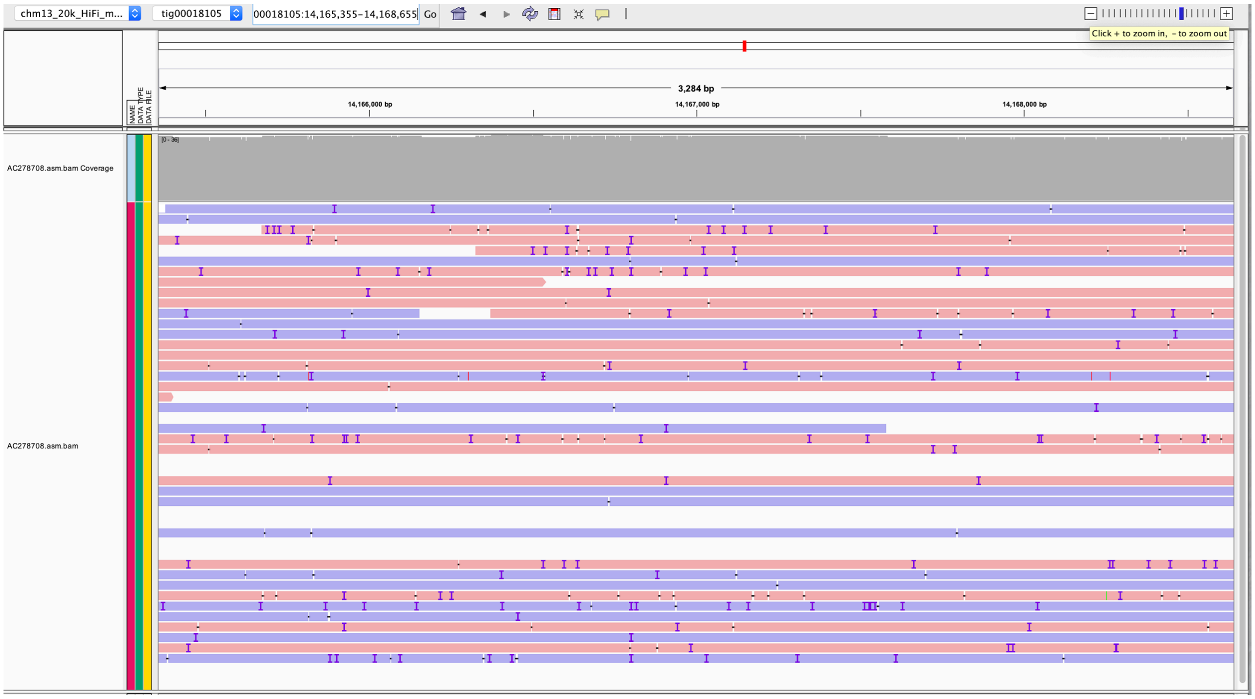


Alignments of HiFi reads to the corresponding assembly region, differing from the BAC sequence. No apparent SVs, SNVs, or coverage anomalies are visible.. Read alignments to the BAC sequence are not shown as the breakpoint region is at the end of the BAC and thus has a coverage drop due to edge effects. The HiCanu assembled version is further supported by GRCh38.p13.

**Supplementary Figure 5. Evaluation of BAC AC278985.1.**

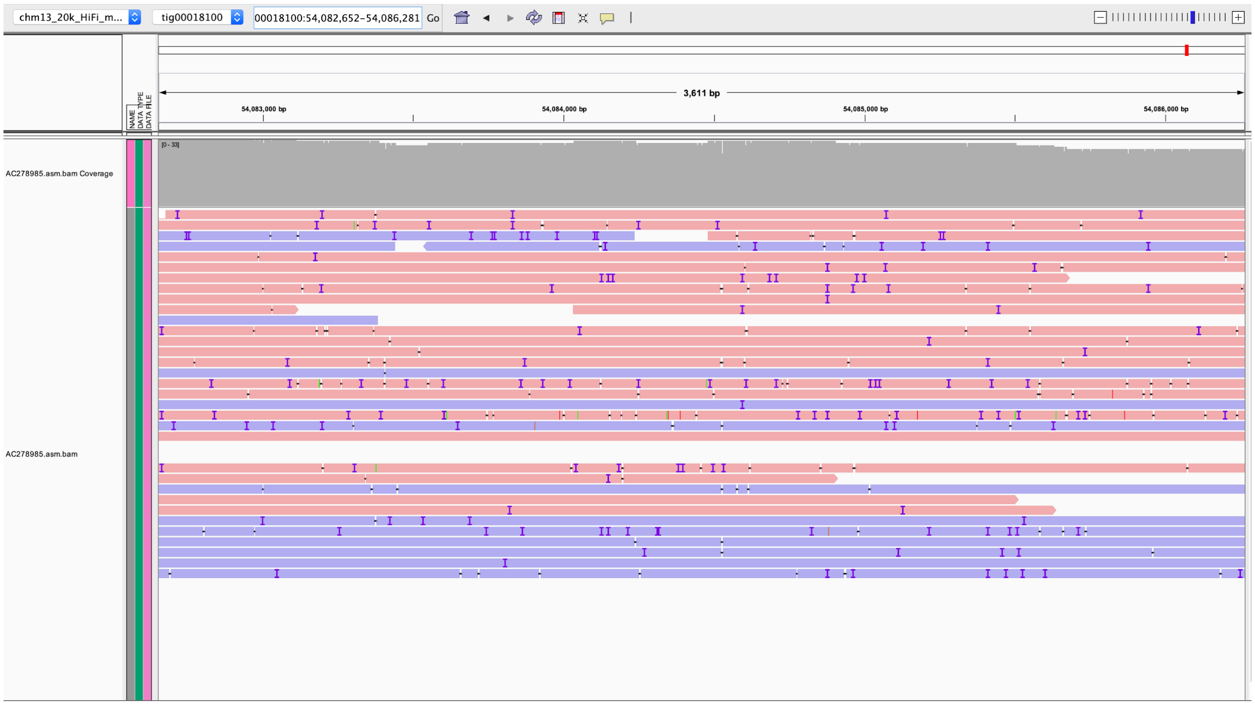


Alignments of HiFi reads to the assembly region, differing from the BAC sequence. No apparent SVs, SNV, or coverage anomalies are visible. Read alignments to the BAC sequence are not show as the breakpoint region is at the end of the BAC and thus has a coverage drop due to edge effects. The HiCanu assembled version is further supported by GCRh38.p13.

**Supplementary Figure 6. Unresolved repeats by class in CHM13 assemblies.**

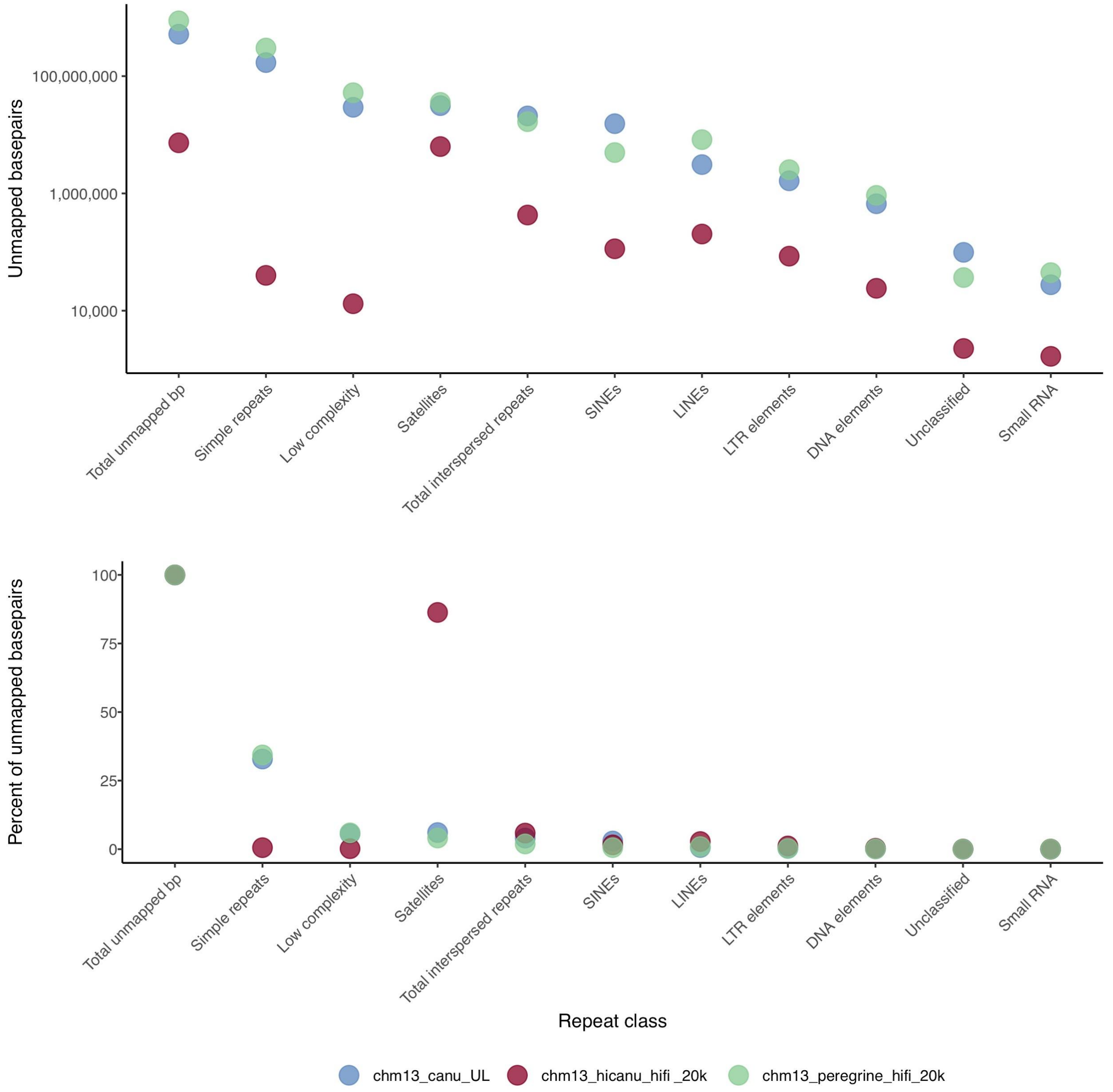


The total unmapped bases (top, log-scaled) and fraction of unmapped bases (bottom) in three assemblies of CHM13. Reads were aligned to the assembly with minimap2 v2.17 (`minimap2 -ax asm20 --secondary=no -s 4000 {input.reads} | samtools view -b - > {output.bam}`), converted to fasta (`samtools fasta -f 4 {output.bam} > {output.fasta}`), and repeats identified by RepeatMasker (`RepeatMasker -species human -e wublast -dir  {ouput.dir} {output.fasta}`). HiCanu (red) improves over both peregrine (green) and ONT UL (blue) assemblies in all repeat types, most notably in simple repeats and low complexity sequences. The majority of remaining unrepresented sequences (over 80% of total) in HiCanu are satellite repeats.

**Supplementary Figure 7. Predicted human assembly continuity based on sequencing read length and accuracy.**

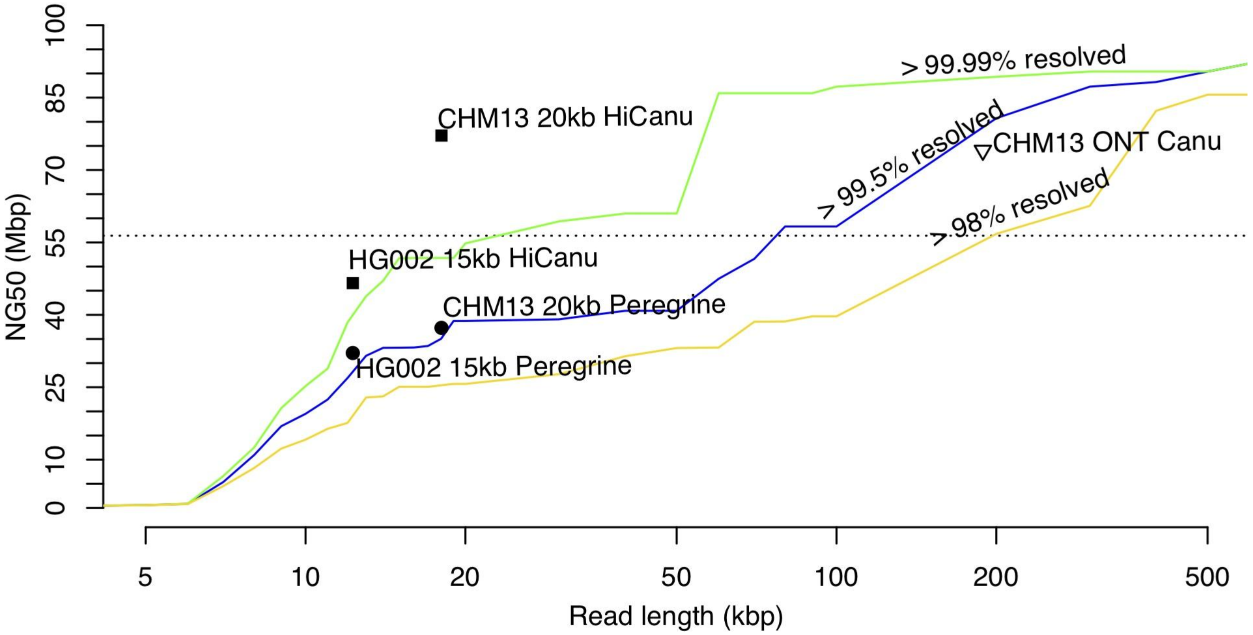


Assembly continuity (NG50 contig size) is dependent on read length (x-axis) and read accuracy (colored curves). Model curves are plotted for three hypothetical assemblers able to separate repeats with 98%, 99.5%, and 99.99% average sequence identity. Points represent real assemblies from Peregrine HiFi (Chin and Khalak 2019), HiCanu HiFi (this work), and Canu Nanopore (Miga et al. 2019). The dashed line corresponds to the continuity of human reference genome GRCh38 (Schneider et al. 2017). Real assemblies typically perform better than predicted by the model, due to the unknown divergence level of many large human repeats which are considered as perfect repeats within the model. In reality, many of the repeat instances appear to have enough variants to be effectively resolved by the assembler.

#### Supplementary Figure 8. HiFi mapping coverage of HiCanu centromere assemblies

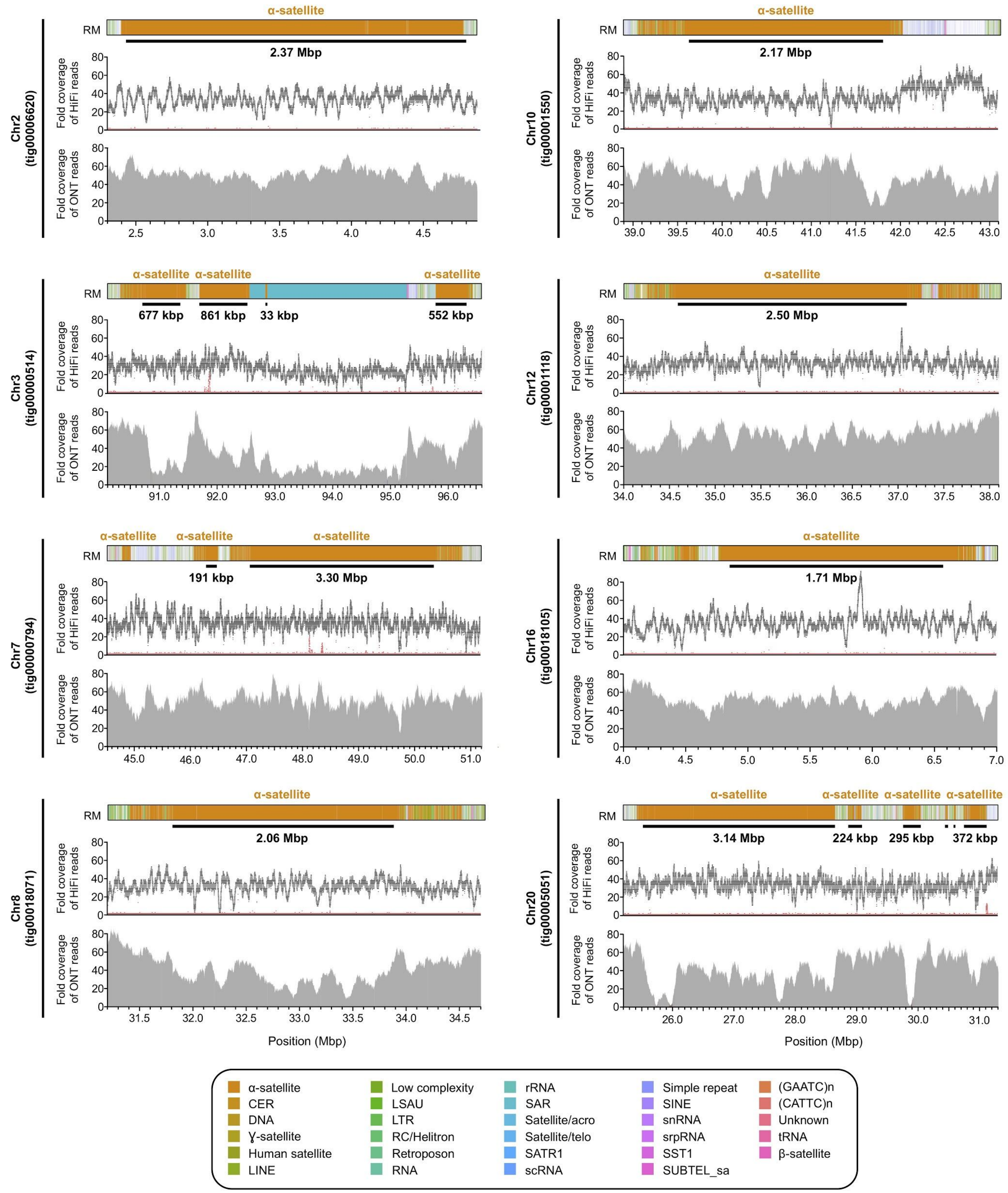


In addition to chromosome 19, HiCanu produced draft assemblies for eight other centromeric satellite arrays of the CHM13 genome. RepeatMasker (RM) annotation reveals the location of α-satellite HOR arrays (marked with a black bar) in each contig. Alignment of HiFi data to each contig reveals even coverage, except for rare dips and spikes in coverage within contigs from chromosomes 8, 10, 12, and 16, which may indicate mis-mapping, mis-assembly, and/or collapse in sequence. Further validation is required to determine the accuracy of these centromeric assemblies.

**Supplementary Figure 9. Sequence identity between the HiCanu D19Z1 or D19Z3 α-satellite arrays and the D19Z1 or D19Z3 α-satellite sequences.**

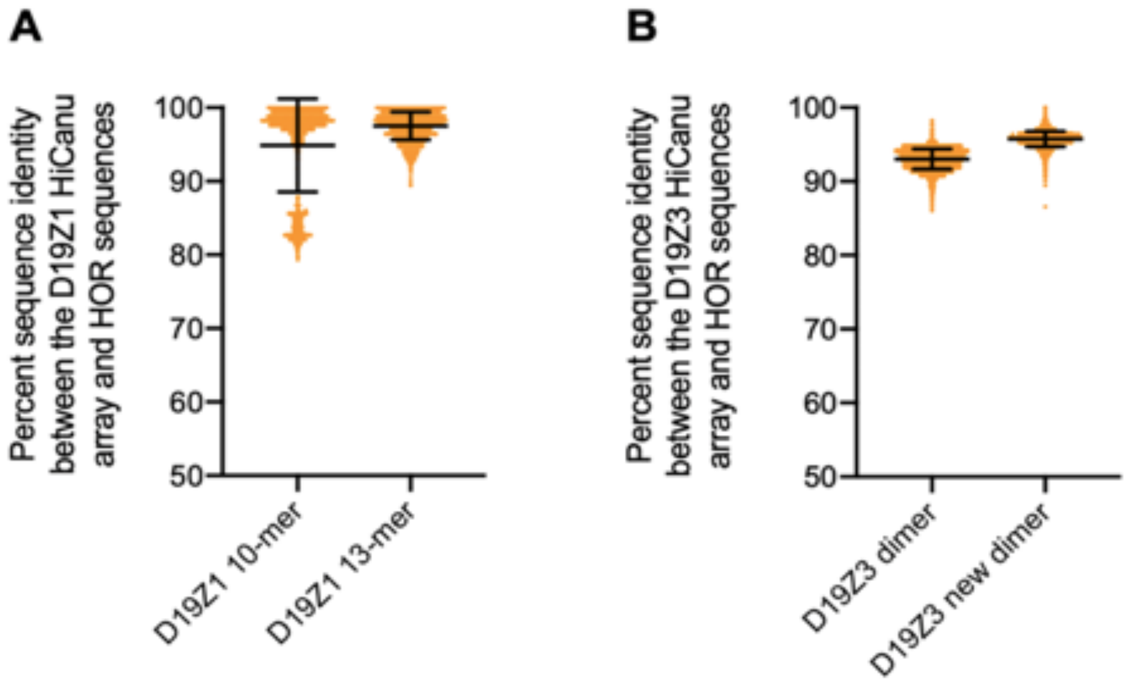


A) Plot showing the sequence identity of each α-satellite repeat within the HiCanu D19Z1 array and the previously identified D19Z1 10-mer (left; Hulsebos et al. 1988; Puechberty et al. 1999; pG-A16, AJ295045.1) or the newly identified D19Z1 13-mer (right). The cluster of low-sequence-identity α-satellite repeats on the left is a result of poor sequence alignment to the 10-mer. When the three newly-identified α-satellite repeats are included to form the 13-mer, this cluster has increased sequence identity, and the overall mean sequence identity increases (94.87 +/- 6.32% vs. 97.51 +/- 1.91%). B) Plot showing the sequence identity of each α-satellite repeat within the HiCanu D19Z3 array and the previously identified D19Z3 dimer (left; Baldini et al. 1989; pC1.8, M26919 and M26920) or the newly identified D19Z3 dimer (right). Almost all α-satellite repeats within the HiCanu D19Z3 array have higher sequence identity to the newly identified dimer than to the previously published dimer (95.74 +/- 1.05% vs. 93.02 +/- 1.37%). Mean +/- SD is shown.

**Supplementary Figure 10. Chromosome 19 D19Z1, D19Z2, and D19Z3 α-satellite HOR array structures.**

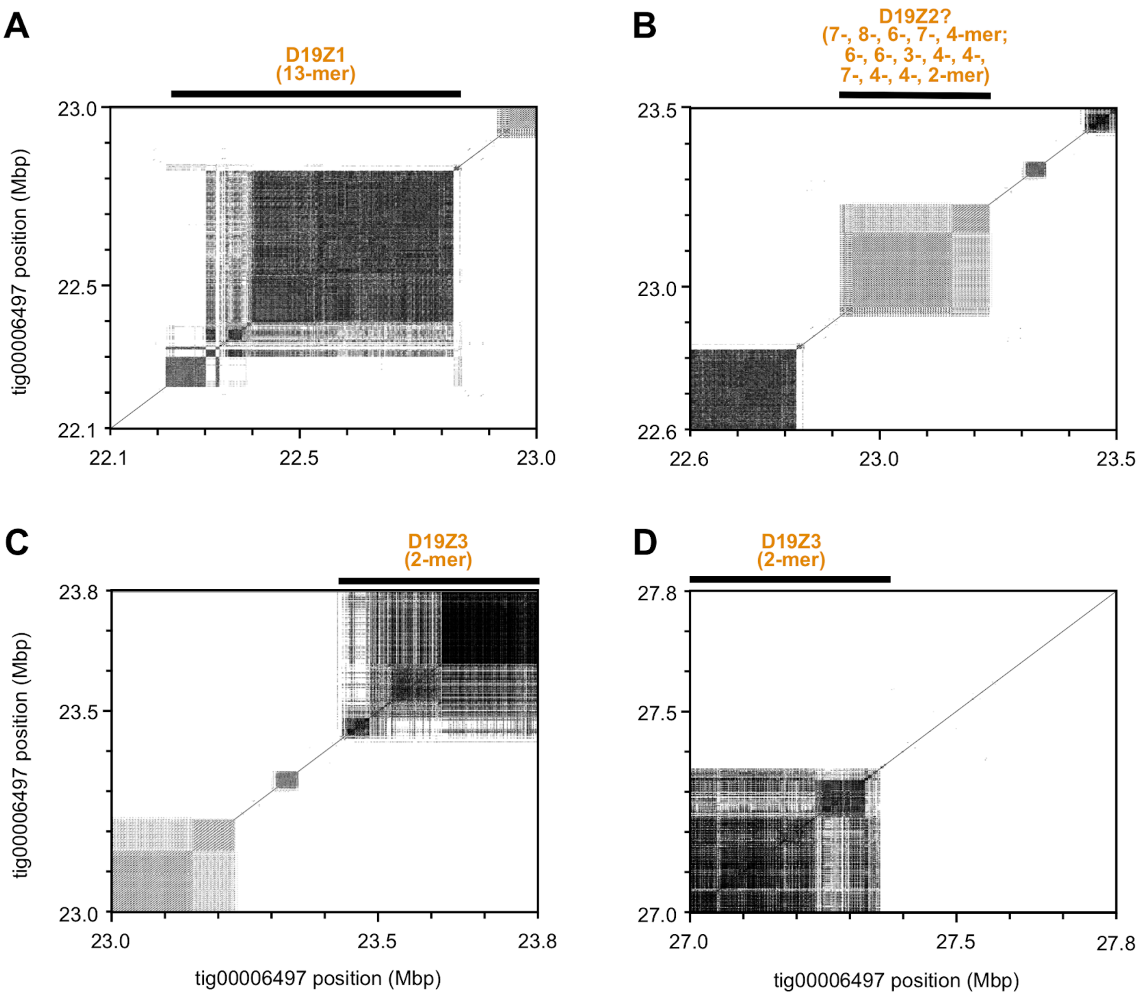


Dot plots showing the high sequence identity and structure of the D19Z1 array (A), D19Z2 array (B), and edges of the D19Z3 array (C, D). The D19Z2 array has two complex HOR structures that can be observed in panel B (see Main Text for details). Because the HOR array structures in this region do not match the expected pG-A16 repeat structure, we designate it here as 'D19Z2?'. Dot plots were generated with a word length of 100.

**Supplementary Figure 11. Segmental duplications associated with contig ends.**

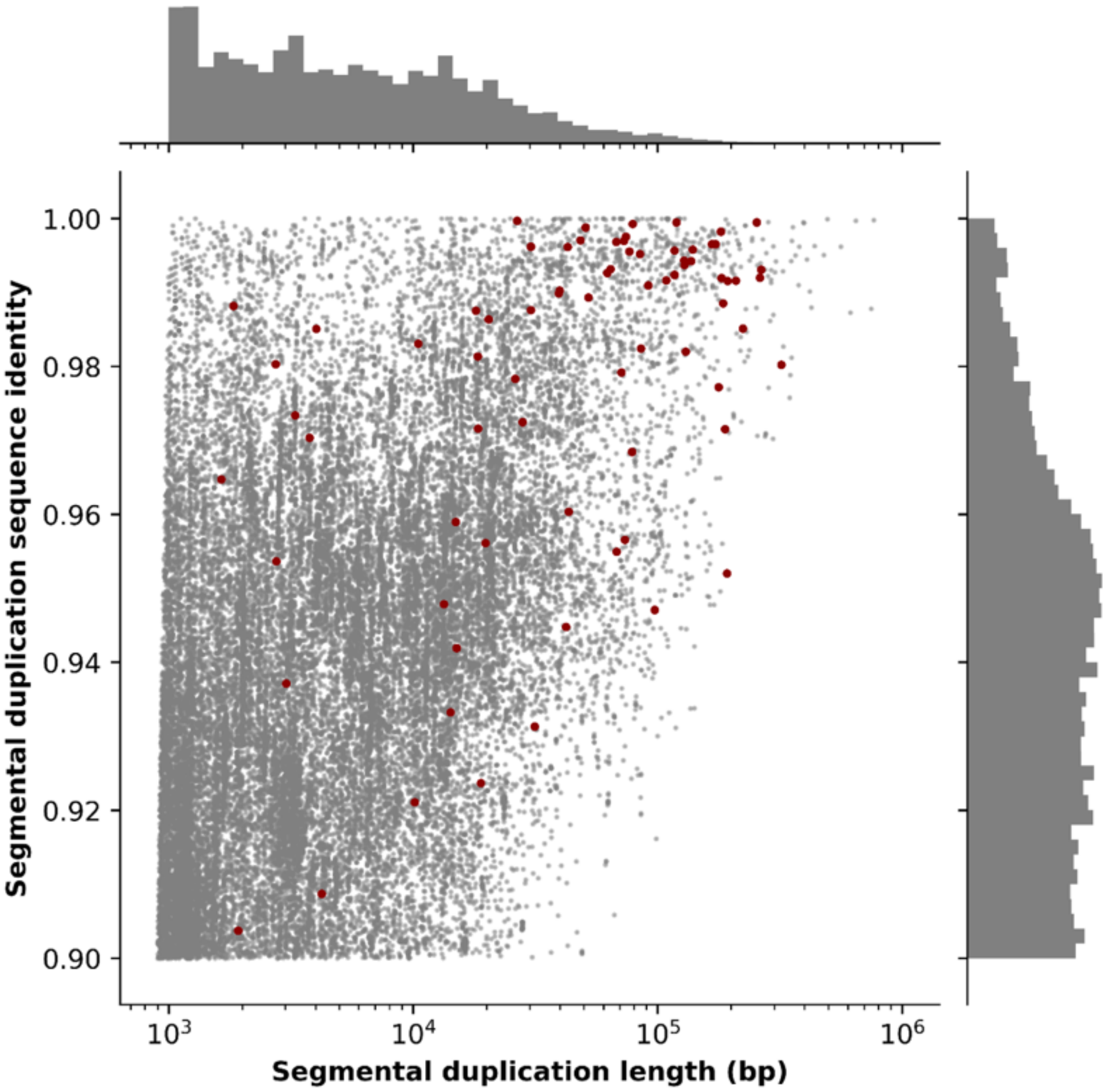


Shown are the sequence identity and length of all segmental duplications in GRCh38 (gray). Red data points indicate the largest segmental duplication within 10 kbp of the 95 contig ends located within segmental duplications in the CHM13 20 kbp HiCanu assembly.

**Supplementary Figure 12. Coverage drop in defensin reference.**

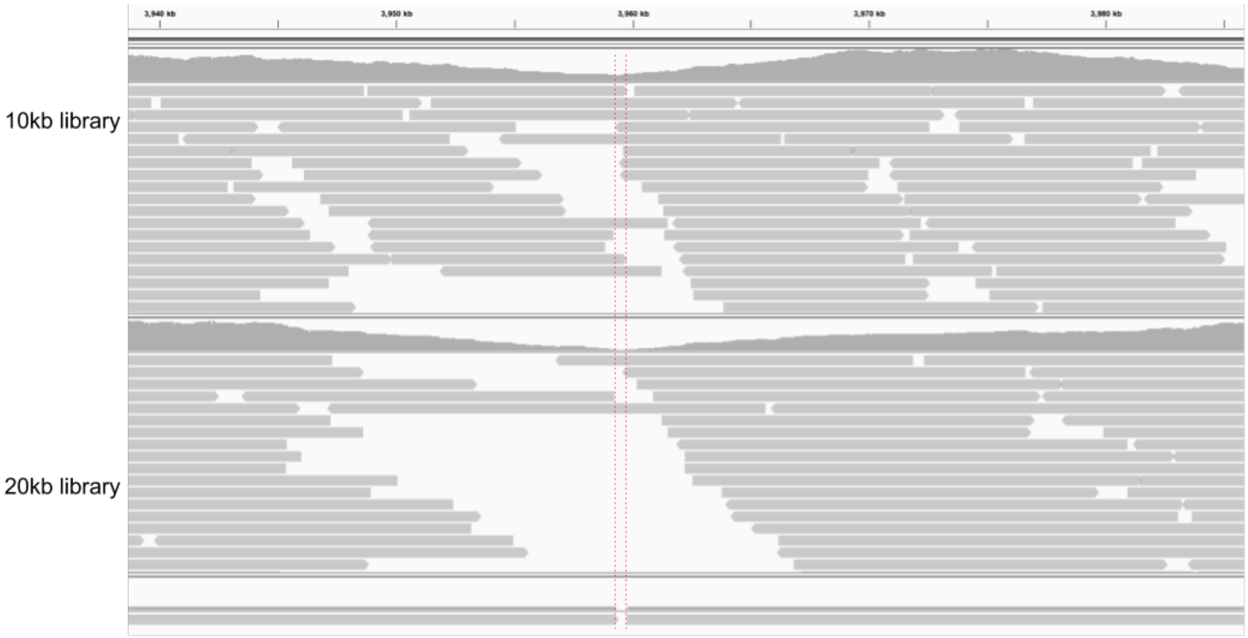


Contig ‘break’ in the HiCanu 20 kbp assembly (highlighted in red) corresponds to a coverage drop in both the 10 kbp and 20 kbp HiFi library. The region immediately upstream of the gap contains a >200bp (AAAGG) simple-sequence repeat.

**Supplementary Figure 13. Bubble contig analysis in HiCanu.**

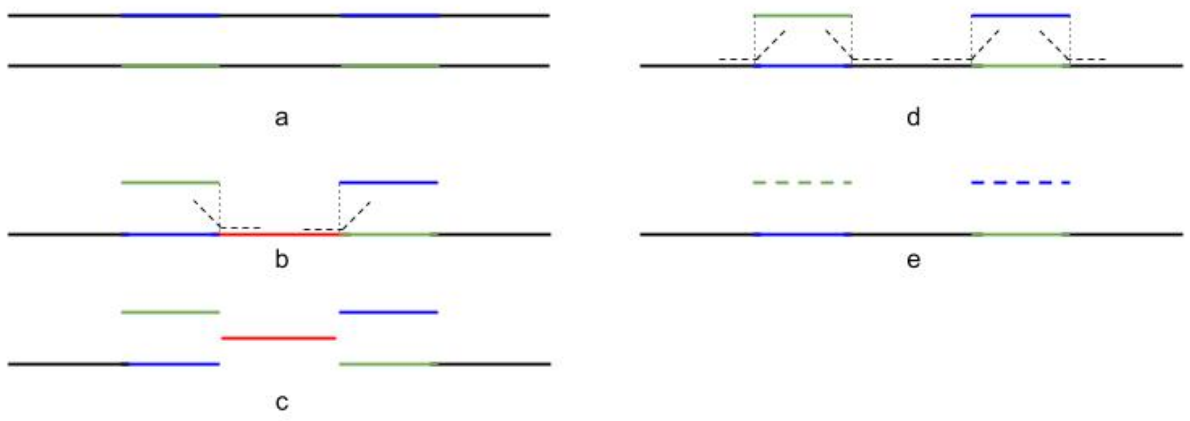


**a.** Genomic area, containing two regions of relatively high heterozygosity (blue/green) divided by a long homozygous region (black). **b.** The homozygous region is typically classified as a genomic repeat due to the reads at boundaries between homozygous and flanking heterozygous regions. **c.** The large pseudo-haplotype contig is broken at the region boundaries. **d.** In HiCanu, bubble contigs (see main text for the definition) are first detected and the reads crossing the regions boundary are no longer considered in repeat detection. **e.** The pseudo-haplotype contig is not split, leaving a continuous pseudo-haplotype.

### Supplementary Tables

**Supplementary Table 1. Alignment-based read identity stats.**

| | **raw reads** | **compressed** | **OEA-corrected** | **Microsatellite mask** |
|---|---|---|---|---|
| Total | 262 664 | 263 526 | 263 529 | 263 559 |
| =100% identity | 907 | 93 840 | 206 711 | 256 259 |
| <100% identity | 261 757 | 169 686 | 56 818 | 7 300 |
| Percent with 100% identity alignment | 0.35% | 35.61% | 78.44% | 97.23% |

The analysis used 20 kbp HiFi CHM13 library. All reads were aligned to the recently finished chrX reference. Reads were output from HiCanu after compression, after OEA-correction, and ignoring errors within microsatellite regions.

**Supplementary Table 2. BUSCO statistics for *D. melanogaster* assemblies.**

| Assembly | Size (Mbp) | NG50 (Mbp) | Complete BUSCOs | Duplicated BUSCOs |
|---|---|---|---|---|
| Canu CLR | 293.73 | 15.20 | 98.8% | 84.7% |
| Peregrine | 162.41 | 12.68 | 98.1% | 1.1% |
| Canu | 292.83 | 13.72 | 99.0% | 92.0% |
| HiCanu | 300.93 | 20.16 | 98.9% | 94.6% |

A genome size of 143,726,002 was used for NG50 computation. Only contigs >=50 kbp were included for analysis.

**Supplementary Table 3. QUAST results for human assemblies.**

| Genome | Assembly | NGA50 (Mbp) | # diffs | # diffs outside known SVs/centromere/segdups |
|---|---|---|---|---|
| CHM13 | Canu ONT | **23.83** | 5,179 | **105** |
| | Canu | 18.96 | 14,501 | 123 |
| | Peregrine | 21.59 | 3,017 | 117 |
| | HiCanu | 21.72 | 15,148 | 140 |
| HG00733 | Canu ONT | 15.92 | 2,632 | 198 |
| | Haplotype asm hap1 | 15.77 | 3,551 | 165 |
| | Haplotype asm hap2 | 15.77 | 3,463 | 143 |
| | Canu | 15.14 | 22,144 | 252 |
| | Peregrine | **21.93** | 3,416 | 215 |
| | HiCanu | 20.51 | 27,721 | 304 |
| | HiCanu (primary) | 16.85 | 13,787 | **93** |
| HG002 | Canu ONT | 15.79 | 2,320 | 171 |
| | Haplotype asm hap1 | 14.33 | 5,854 | 355 |
| | Haplotype asm hap2 | 13.50 | 5,370 | 343 |
| | Canu | 17.01 | 15,120 | 156 |
| | Peregrine | 16.52 | 2,993 | 102 |
| | HiCanu | **21.72** | 21,672 | 229 |
| | HiCanu (primary) | 17.65 | 12,019 | **73** |

NGA50s were computed using the following size: 3,098,794,149. GRCh38 excluding unassigned and alts contigs was used as a reference. Structural variants and identity computed using QUAST v5.0.2. “# diffse” is the sum of relocations, translocations, and inversions. Using scripts from Safin et al., we ignored differences within the regions with known variants (for HG002 only) and regions in centromeres and segmental duplications since these regions are likely enriched for real differences between the reference and sequenced genomes. Haplotype-resolved assemblies are from Garg et al. 2019 (HG002) and Porubsky et al. 2019 (HG00733)

**Supplementary Table 4: CHM13 10 kbp HiFi assemblies**

| Assembly | NG50 Mbp | Total Gbp | Illumina QV | BACs | BAC QV | NGA50 (Mbp) | # QUAST diffs | # diffs outside contigs <50kb, known SVs/centromere/seg dups | Complete BUSCOs |
|---|---|---|---|---|---|---|---|---|---|
| Canu + Racon x2 | 29.1 | 2.94 | **61.8** | 155/341 | 36.9 | 17.39 | 6,809 | 112 | 94.8% |
| Peregrine | 29.7 | 2.83 | 51.3 | 122/341 | 34.8 | 16.87 | **1,219** | **97** | 94.7% |
| HiCanu | **39.3** | 3.03 | **61.7** | **309/341** | **40.8** | **19.24** | 12,001 | 114 | **95.1%** |

CHM13 10 kbp assemblies from Canu, Peregrine, and HiCanu were evaluated as before. All statistics are reported on contigs >= 50 kbp.

**Supplementary Table 5. CHM13 corrected BACs results.**

| Genome | Assembly | Original BACs resolved | Corrected BACs resolved |
|---|---|---|---|
| CHM13 | Canu ONT (Miga et al. 2019) | 314/341 | 323/341 |
| | Canu 10kb | 155/341 | 157/341 |
| | Canu 20kb | 308/341 | 316/341 |
| | Peregrine 10kb | 122/341 | 124/341 |
| | Peregrine 20kb | 136/341 | 137/341 |
| | HiCanu 10kb | 309/341 | 320/341 |
| | HiCanu 20kb | 326/341 | **337/341** |

CHM13 BACs were downloaded from the library VMRC59 at NCBI. BACs putatively identified as incorrect were replaced by their reconstruction from the HiCanu 20 kbp assembly. Statistics were computed using the scripts at https://github.com/skoren/bacValidation. Despite the BAC corrections coming from the assembly of the 20kbp HiFi dataset, 8 were resolved by at least two other assemblies (Peregrine 10 kbp/20 kbp, Canu 10 kbp/20 kbp/Canu UL), and three were resolved in one other assembly (AC278859.1 Canu ONT, AC278985.1 Canu ONT, AC279108.1 Canu 10 kbp), supporting the updated sequences.

**Supplementary Table 6. CHM13 collapsed bases.**

| Datatype | Assembly | Collapsed (Mbp) | Expanded (Mbp) |
|---|---|---:|---:|
| ONT UL | Canu ONT | 35.00 | 254.36 |
| HiFi 10kbp | Canu + Racon x2 | 53.74 | 183.60 |
| | Peregrine | 45.57 | 328.42 |
| | HiCanu | 27.41 | 78.56 |
| HiFi 20kbp | Canu | 24.88 | 71.03 |
| | Peregrine | 44.85 | 309.27 |
| | HiCanu | **22.80** | **56.73** |

CHM13 collapses were evaluated using SDA as in Vollger et al. 2019. All contigs in the assembly are included for analysis. The *collapsed* bases are the assembly bases with “higher than expected coverage, while the *expanded* bases is the estimate of how much sequence is contained in the collapsed regions, had they been correctly assembled.

**Supplementary Table 7. BUSCO and phase block statistics.**

| Genome | Assembly | Complete BUSCOs | Duplicated BUSCOs | Phase block NG50 (Mbp) | Intra-block switch rate |
|---|---|---|---|---|---|
| CHM13 | Canu ONT | 79.3% | 1.0% | N/A | N/A |
| | Canu | 94.7% | 1.5% | N/A | N/A |
| | Peregrine | 94.8% | 1.4% | N/A | N/A |
| | HiCanu | 94.1% | 2.7% | N/A | N/A |
| HG00733 | Canu ONT (alts) | 66.3% (0.3%) | 0.8% (0.0%) | 0.23 (0.00) | 18.27 (11.74) |
| | Haplotype asm hap1 (hap2)[a] | 94.8% (95.2%) | 1.4% (1.4%) | 5.14 (4.49) | 0.43 (0.56) |
| | Canu primary (alts) | 94.8% (22.7%) | 1.5% (1.2%) | 0.17 (0.00) | 5.20 (0.12) |
| | Peregrine (alts) | 95.0% (0.5%) | 1.3% (0.1%) | 0.15 (0.00) | 8.47 (5.01) |
| | HiCanu primary (alts) | 94.9% (77.2%) | 1.4% (1.9%) | 0.62 (0.14) | 0.45 (0.12) |
| HG002 | Canu ONT (alts) | 63.4% (0.2%) | 0.8% (0.0%) | 0.23 (0.00) | 11.24 (0.97) |
| | Haplotype asm hap1 (hap2)[b] | 94.7% (95.0%) | 1.2% (1.2%) | 12.82 (11.42) | 0.13 (0.13) |
| | Canu primary (alts) | 92.1% (5.5%) | 1.2% (0.3%) | 0.15 (0.00) | 3.18 (0.07) |
| | Peregrine (alts) | 95.1% (0.6%) | 1.2% (0.1%) | 0.15 (0.00) | 5.43 (0.30) |
| | HiCanu primary (alts) | 94.8% (77.1%) | 1.2% (1.7%) | 0.65 (0.10) | 0.24 (0.01) |

All contigs were included for analysis, regardless of length. [a]Assembly generated in Porubsky et al. 2019. [b]Assembly generated in Garg et al. 2019.

**Supplementary Table 8. HLA gene truth typing from Chin et al. 2019 (HG002) and Shafin et al. 2019 (HG00733).**

| Genome | Gene | Expected haplotype1 | Expected haplotype2 |
|---|---|---|---|
| HG00733 | HLA-A | 30:02:01G | 24:02:01G |
| | HLA-B | 18:01:01G | 35:02:01G |
| | HLA-C | 05:01:01G | 04:01:01G |
| | HLA-DQA1 | 05:01:01G | 05:01:01G |
| | HLA-DQB1 | 02:01:01G | 03:01:01G |
| | HLA-DRB1 | 03:01:01G | 11:04:01G |
| HG002 | HLA-A | 26:01:01G | 01:01:01G |
| | HLA-B | 38:01:01G | 35:08:01G |
| | HLA-C | 12:03:01G | 04:01:01G |
| | HLA-DQA1 | 03:01:01G | 01:01:01G |
| | HLA-DQB1 | 03:02:01G | 05:01:01G |
| | HLA-DRB1 | 04:02:01G | 10:01:01G |

**Supplementary Table 9. HLA gene typing results across different assemblies.**

| Genome | Assembly | Contig | Gene | Called | Edit Distance |
|---|---|---|---|---|---|
| HG0733 | Canu | tig00032730 | HLA-A | A*30:02:01G | 4 |
| | | tig00032730 | HLA-B | B*35:02:01G | 0 |
| | | tig00032730 | HLA-C | C*04:01:01G | 0 |
| | | tig00062315 | HLA-DQA1 | DQA1*05:01:01G | 0 |
| | | tig00062315 | HLA-DQB1 | DQB1*03:01:01G | 0 |
| | | tig00062315 | HLA-DRB1 | DRB1*11:04:01G | 0 |
| | | tig00037512 | HLA-A | A*24:02:01G | 0 |
| | | tig00037527 | HLA-B | B*18:01:01G | 0 |
| | | tig00037527 | HLA-C | C*05:01:01G | 0 |
| | | tig00038945 | HLA-DQA1 | DQA1*05:01:01G | 0 |
| | | tig00038945 | HLA-DQB1 | DQB1*02:01:01G | 0 |
| | | tig00038955 | HLA-DRB1 | DRB1*03:01:01G | 0 |
| | HiCanu | tig00018936 | HLA-A | A*30:02:01G | 0 |
| | | tig00018936 | HLA-B | B*35:02:01G | 0 |
| | | tig00018936 | HLA-C | C*04:01:01G | 0 |
| | | tig00023025 | HLA-DQA1 | DQA1*05:01:01G | 0 |
| | | tig00023025 | HLA-DQB1 | DQB1*03:01:01G | 0 |
| | | tig00023025 | HLA-DRB1 | DRB1*11:04:01G | 0 |

| | | | | | |
|---|---|---|---|---|---|
| | | tig00029466 | HLA-A | A*24:02:01G | 0 |
| | | tig00029414 | HLA-B | B*18:01:01G | 0 |
| | | tig00029414 | HLA-C | C*05:01:01G | 0 |
| | | tig00023598 | HLA-DQA1 | DQA1*05:01:01G | 0 |
| | | tig00023598 | HLA-DQB1 | DQB1*02:01:01G | 0 |
| | | tig00023598 | HLA-DRB1 | DRB1*03:01:01G | 0 |
| | StrandSeq HiFi | 000030F | HLA-A | A*30:02:01G | 0 |
| | | 000030F | HLA-B | B*35:02:01G | 0 |
| | | 000030F | HLA-C | C*04:01:01G | 0 |
| | | 000030F | HLA-DQA1 | DQA1*05:01:01G | 0 |
| | | 000030F | HLA-DQB1 | DQB1*03:01:01G | 0 |
| | | 000030F | HLA-DRB1 | DRB1*11:04:01G | 0 |
| | | 000118F | HLA-A | A*30:02:01G | 0 |
| | | 000118F | HLA-B | B*35:02:01G | 0 |
| | | 000118F | HLA-C | C*05:01:01G | 0 |
| | | 000031F | HLA-DQA1 | DQA1*05:01:01G | 0 |
| | | 000031F | HLA-DQB1 | DQB1*02:01:01G | 0 |
| | | 000031F | HLA-DRB1 | DRB1*03:01:01G | 0 |
| | Peregrine | 000029F | HLA-A | A*30:02:01G | 4 |
| | | 000029F | HLA-B | B*35:02:01G | 1 |

| | | | | | |
|---|---|---|---|---|---|
| | | 000029F | HLA-C | C*04:103 | 3 |
| | | 000029F | HLA-DQA1 | DQA1*05:01:01G | 0 |
| | | 000029F | HLA-DQB1 | DQB1*03:01:01G | 0 |
| | | 000029F | HLA-DRB1 | DRB1*11:04:01G | 0 |
| HG002 | Canu | tig00038929 | HLA-A | A*01:01:01G | 0 |
| | | tig00038929 | HLA-B | B*38:01:01G | 0 |
| | | tig00038929 | HLA-C | C*04:01:01G | 0 |
| | | tig00038927 | HLA-DQA1 | DQA1*03:01:01G | 0 |
| | | tig00038927 | HLA-DQB1 | DQB1*03:02:01G | 0 |
| | | tig00038927 | HLA-DRB1 | DRB1*04:02:01 | 0 |
| | | tig00013892 | HLA-A | A*26:01:01G | 0 |
| | | tig00013881 | HLA-B | B*35:08:01G | 0 |
| | | tig00013903 | HLA-C | C*12:03:01G | 0 |
| | | tig00007817 | HLA-DQA1 | DQA1*01:01:01G | 0 |
| | | tig00007817 | HLA-DQB1 | DQB1*05:01:01G | 0 |
| | | tig00007817 | HLA-DRB1 | DRB1*10:01:01G | 0 |
| | HiCanu | tig00009386 | HLA-A | A*01:01:01G | 0 |
| | | tig00009386 | HLA-B | B*35:08:01G | 0 |
| | | tig00009386 | HLA-C | C*04:01:01G | 0 |
| | | tig00009386 | HLA-DQA1 | DQA1*03:01:01G | 0 |

| | | | | | |
|---|---|---|---|---|---|
| | | tig00009386 | HLA-DQB1 | DQB1*03:02:01G | 0 |
| | | tig00009386 | HLA-DRB1 | DRB1*04:02:01 | 0 |
| | | tig00009431 | HLA-A | A*26:01:01G | 0 |
| | | tig00017533 | HLA-B | B*38:01:01G | 0 |
| | | tig00017533 | HLA-C | C*12:03:01G | 0 |
| | | tig00009488 | HLA-DQA1 | DQA1*01:01:01G | 0 |
| | | tig00009488 | HLA-DQB1 | DQB1*05:01:01G | 0 |
| | | tig00009488 | HLA-DRB1 | DRB1*10:01:01G | 0 |
| | TrioCanu HiFi | tig00004595\|arrow\|arrow | HLA-A | A*01:01:01G | 0 |
| | | tig00004595\|arrow\|arrow | HLA-B | B*35:08:01G | 0 |
| | | tig00004595\|arrow\|arrow | HLA-C | C*04:01:01G | 0 |
| | | tig00004595\|arrow\|arrow | HLA-DQA1 | DQA1*01:01:01G | 0 |
| | | tig00004595\|arrow\|arrow | HLA-DQB1 | DQB1*05:01:01G | 0 |
| | | tig00004595\|arrow\|arrow | HLA-DRB1 | DRB1*10:01:01G | 0 |
| | | tig00004691\|arrow\|arrow | HLA-A | A*26:01:01G | 0 |
| | | tig00004691\|arrow\|arrow | HLA-B | B*38:01:01G | 0 |
| | | tig00004691\|arrow\|arrow | HLA-C | C*12:03:01G | 0 |
| | | tig00004302\|arrow\|arrow | HLA-DQA1 | DQA1*03:01:01G | 0 |
| | | tig00004302\|arrow\|arrow | HLA-DQB1 | DQB1*03:02:01G | 0 |
| | | tig00004302\|arrow\|arrow | HLA-DRB1 | DRB1*04:02:01 | 0 |

| | | | | | |
|---|---|---|---|---|---|
| | | HG002-S16-H1-000005F | HLA-A | A*01:01:01G | 0 |
| | | HG002-S16-H1-000005F | HLA-B | B*35:08:01G | 0 |
| | | HG002-S16-H1-000005F | HLA-C | C*04:01:01G | 0 |
| | | HG002-S16-H1-000003F | HLA-DQA1 | DQA1*01:01:01G | 0 |
| | | HG002-S16-H1-000003F | HLA-DQB1 | DQB1*05:01:01G | 0 |
| | | HG002-S16-H1-000003F | HLA-DRB1 | DRB1*03:96<br>DRB1*13:178 | 46 |
| | | HG002-S16-H2-000005F | HLA-A | A*26:01:01G | 0 |
| | | HG002-S16-H2-000005F | HLA-B | B*38:01:01G | 0 |
| | | HG002-S16-H2-000005F | HLA-C | C*12:03:01G | 0 |
| | | HG002-S16-H2-000002F | HLA-DQA1 | DQA1*03:01:01G | 0 |
| | | HG002-S16-H2-000002F | HLA-DQB1 | DQB1*03:02:01G | 0 |
| | Hi-C HiFi | HG002-S16-H2-000002F | HLA-DRB1 | DRB1*04:02:01<br>DRB1*04:02:05 | 1 |
| | | 000029F | HLA-A | A*01:01:01G | 0 |
| | | 000029F | HLA-B | B*38:01:01G | 0 |
| | | 000029F | HLA-C | C*12:03:01G | 0 |
| | | 000029F | HLA-DQA1 | DQA1*03:01:01G | 0 |
| | | 000029F | HLA-DQB1 | DQB1*03:02:01G | 0 |
| | Peregrine | 000029F | HLA-DRB1 | DRB1*04:02:01 | 0 |

Results for Peregrine, Canu and HiCanu assemblies (this paper), as well as results of a previous HiFi TrioCanu assembly (Wenger et al. 2019) and recent Hi-C (Garg et al. 2019) and StrandSeq (Porubsky et al. 2019) assemblies. The trio binning results are accurate and in phase across the entire MHC, capturing the locus in two contigs (one per haplotype). HiCanu assemblies include all expected alleles (without errors), but the reconstruction is more fragmented with a haplotype switch in the primary contig set between class I and class II genes in HG002 and after HLA-A in HG00733. The Hi-C based assembly from Garg et al. is in phase but has two non-0 edit distance genes, one of which matches neither haplotype. The StrandSeq phased assembly from Porubsky et al. only captures one version of the HLA-A and HLA-B genes, losing the alleles from other haplotype.

**Supplementary Table 10. CHM13 centromeres identified by RepeatMasker**

| Tig ID | Chromosome | Length (Mbp) | Centromere Start (Mbp) | Centromere End (Mbp) |
|---|---|---:|---:|---:|
| tig00006620 | chr2 | 49.17 | 2.3 | 4.9 |
| tig00000514 | chr3 | 199.46 | 90.0 | 96.6 |
| tig00000794 | chr7 | 147.14 | 44.5 | 51.2 |
| tig00018071 | chr8 | 129.87 | 31.2 | 34.7 |
| tig00001550 | chr10 | 112.08 | 38.9 | 43.1 |
| tig00001118 | chr12 | 126.71 | 34.0 | 38.1 |
| tig00018105 | chr16 | 26.11 | 4.0 | 7.0 |
| tig00006497 | chr19 | 46.04 | 21.7 | 28.1 |
| tig00005051 | chr20 | 65.33 | 25.3 | 31.3 |

Positions of internal centromeres (at least 500 kbp away from the contig start/end) identified by RepeatMasker and manually adjusted based on visual inspection of RepeatMasker results.
